# Protein lactylation induced by neural excitation

**DOI:** 10.1101/2021.02.02.428370

**Authors:** Hideo Hagihara, Hirotaka Shoji, Hikari Otabi, Atsushi Toyoda, Kaoru Katoh, Masakazu Namihira, Tsuyoshi Miyakawa

## Abstract

Lactate is known to have diverse roles in the brain at the molecular and behavioral levels under both physiological and pathophysiological conditions, such as learning and memory and regulation of mood. Recently, a novel post-translational modification called lysine lactylation has been found in histone H3 of mouse macrophages, and the lactylation levels paralleled the intracellular lactate levels^1^. However, it is unknown whether lysine lactylation occurs in brain cells, and if it does, whether lactylation is induced by the stimuli that accompany changes in lactate levels. Herein, we reveal that lysine lactylation in brain cells is regulated by systemic changes in lactate levels, neural excitation, and behaviorally relevant stimuli. Lysine lactylation levels were increased by lactate treatment and by high-potassium-induced depolarization in cultured primary neurons; these increases were attenuated by pharmacological inhibition of monocarboxylate transporter 2 and lactate dehydrogenase, respectively, suggesting that both cell-autonomous and non-cell-autonomous neuronal mechanisms are involved in overall lysine lactylation. *In vivo*, electroconvulsive stimulation increased lysine lactylation levels in the prefrontal cortices of mice, and its levels were positively correlated with the expression levels of the neuronal activity marker c-Fos on an individual cell basis. In the social defeat stress model of depression in which brain lactate levels increase, lactylation levels were increased in the prefrontal cortices of the defeated mice, which was accompanied by increased c-Fos expression, decreased social behaviors, and increased anxiety-like behaviors, suggesting that stress-induced neuronal excitation may induce lysine lactylation, thereby affecting mood-related behaviors. Further, we identified 63 candidate lysine-lactylated proteins in the mouse cortex and found that lactylation levels in histone H1 increased in response to defeat stress. This study may open up an avenue for exploration of a novel role of neuronal activity-induced lactate mediated by protein lactylation in the brain.

## Main text

Lactate in the brain has emerged as an energy substrate and a valuable signaling molecule^2^, and its alteration has been implicated in multiple neuropsychiatric disorders. Among various strains/conditions of animal models of neuropsychiatric disorders, we have identified more than 20 out of 65 strains/conditions of mice that show altered lactate levels in whole brain^3,4^. A recent research identified lysine lactylation (Kla) as a novel post-translational modification in histone protein that can be stimulated by lactate^1^. A report by Zhang et al. suggested that histone H3 Kla mediates a metabolic alteration-related lactate regulation of gene expression in mouse bone marrow-derived macrophages^1^. The following reports have described key roles of lactate-induced Kla in macrophages in the transition of macrophages from a proinflammatory state to a reparative state^5^ and the pathogenesis of lung fibrosis^6^. However, it has not been elucidated whether this Kla occurs and is regulated by exogenous lactate treatment or by stimulation that increases endogenous lactate levels in other types of cells/tissues including brain cells.

### Lactate treatment and neuronal excitation increase protein lactylation in the brain

Kla was detected by western blot analysis in mouse brain tissues, with patterns different from those observed in macrophages and peripheral organs (Fig. 1a, Extended Data Fig. 1). With confocal microscopic imaging, we observed Kla signals throughout the cell, including somata and neurites, with a relatively weak signal in the nucleus region in primary cultured neurons expressing the neuronal marker βIII-tubulin (Fig. 1b). These results suggest that Kla modifications occur in various proteins to varying degrees, potentially with neural cell-specific functional roles different from those found in macrophages.

**Figure 1.**
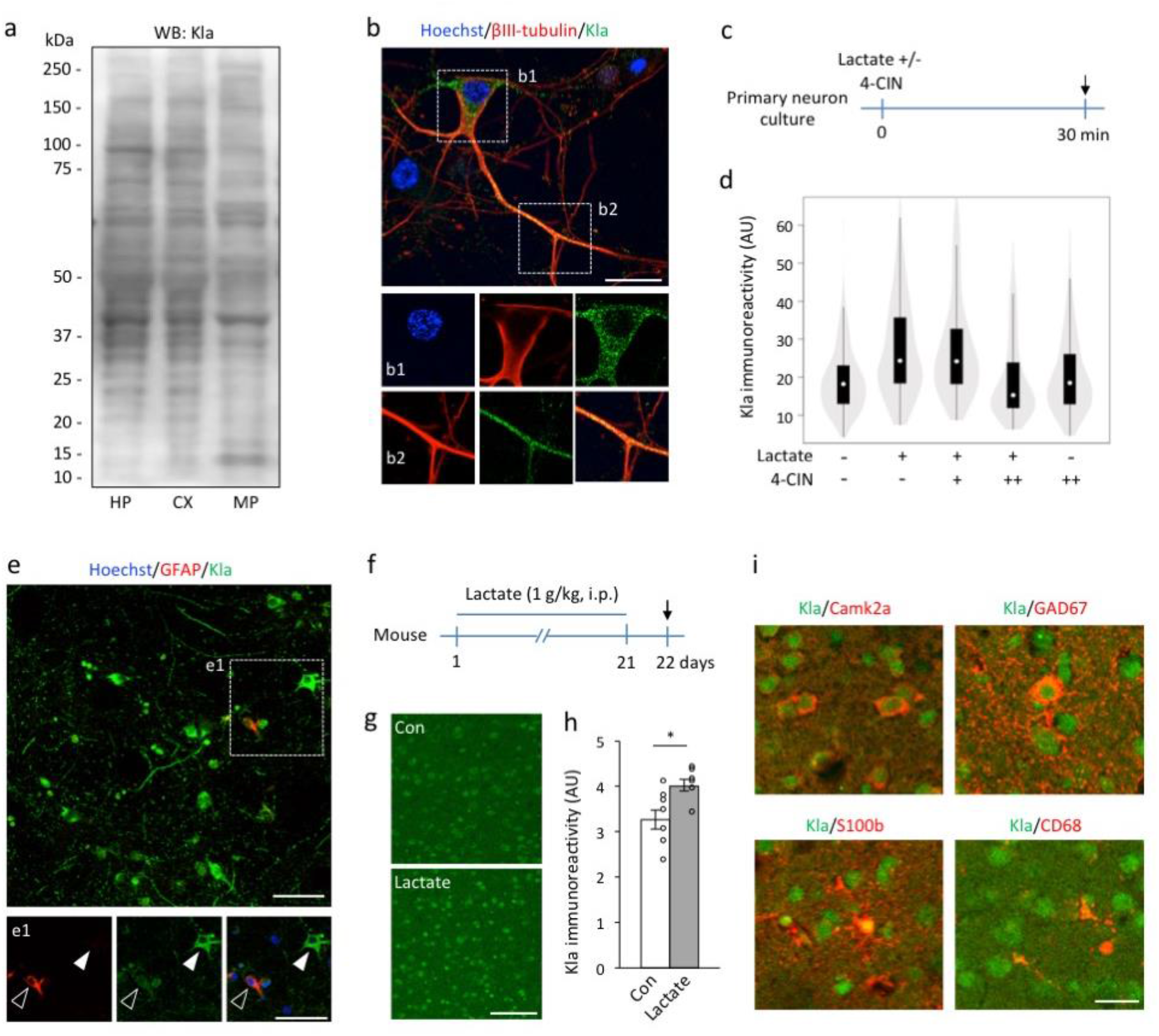
Protein lactylation induced by exogenous lactate in brain cells. (a) Western blot images of lactylated lysine (Kla) in mouse tissues/cells (hippocampus (HIP), cerebral cortex (CX), and resident macrophages (MP). Entire membrane images are provided in the Extended Data Fig. 1. (b) Double immunostaining for Kla and the neuronal marker βIII-tubulin in primary cultured cortical neurons (10 days in vitro, DIV). Scale bar: 20 μm. (c, d) Kla levels in primary cultured cortical neurons (10 DIV) examined 30 minutes after treatment with lactate (25 mM, +) and/or 4-CIN (0.01 mM, +; 1 mM, ++) using immunocytochemistry. Arrow in (c) indicates the sampling time point. At least 200 cells were examined for each condition. Statistical results are provided in Extended Data Table 4. (e) Glial fibrillary acidic protein (GFAP) and Kla double immunostaining at 10 DIV in culture. Open arrows indicate GFAP-positive astrocyte-like cells. Solid arrows indicate GFAP-negative neuronal cells. Scale bar: 50 μm. (f–h) Mice were intraperitoneally injected with 1 g/kg lactate or saline, once daily for 21 days. (g) Kla immunostaining images of the prefrontal cortices (PFCs) of control and lactate-treated mice. Scale bar: 100 μm. (h) Bar graph showing Kla immunoreactivity levels of the PFC of control and lactate-treated mice (P = 0.022, n = 7, 6). (i) Mouse PFC: Double immunostaining images of Kla and the glutamatergic neuron marker Camk2a (upper left panel), the GABAergic neuron marker GAD67 (upper right panel), the astrocyte marker S100b (lower left panel), and the microglial marker CD68 (lower right panel). Scale bar: 25 μm.

We investigated whether extracellular application of lactate stimulates Kla in neuronal cells as seen in macrophages^1^. Lactate treatment significantly increased Kla immunoreactivity in the primary cultured neurons (Fig. 1c, d), as confirmed by an independent experiment performed at a different institute (Extended Data Fig. 2). The increase in Kla immunoreactivity was attenuated in a dose-dependent manner by α-cyano-4-hydroxycinnamate (4-CIN), a monocarboxylate transporter 2 (MCT2) inhibitor that blocks lactate transport into neurons (Fig. 1d). Considering that there are few astrocyte-like cells in the culture (Fig. 1e), lactate derived from astrocytes may minimally affect the lactate contents therein. Furthermore, in an *in vivo* experiment, we found that chronic and systemic treatment with lactate, shown to increase extracellular lactate concentrations in the brain^7^, increased Kla immunoreactivity in the mouse prefrontal cortex (PFC) (Fig. 1f–h), but not in the hippocampus (Extended Data Fig. 3), suggesting that exogenous lactate stimulates protein lactylation in the brain in a region-dependent manner.

To examine in what neural cell types protein lactylation occurs in the brain, we conducted double-immunostaining of Kla with cell type-specific markers on mouse PFC: Camk2a for glutamatergic neurons, GAD67 for GABAergic neurons, S100b for astrocytes, and CD68 for microglia. Kla immunoreactivity was found in all cell types examined, suggesting that protein lactylation may ubiquitously occur in neural cells in the brain (Fig. 1i).

### Neuronal excitation stimulates protein lactylation in the brain

Neuronal excitation increases brain lactate levels^8–12^. We examined lactate and protein lactylation levels using an *in vitro* neuronal excitation model. Primary neuron cultures were treated with a high concentration of potassium ions (high-K) (Fig. 2a) to induce depolarization of neurons *in vitro*. The high-K treatment resulted in increased lactate concentration in the culture supernatant at 30 minutes after the initiation of treatment (Fig. 2b). The intracellular lactate level showed a trend of time-dependent increase and reached significant increase at 2 hours after the initiation of high-K treatment (Fig. 2b). The high-K-induced increase in Kla levels was attenuated by treatment with sodium oxamate (OX), which inhibits lactate dehydrogenase (LDH) activity in neural^13,14^ and other cell types ^1^ (Fig. 2c), suggesting that neuronal excitation may induce protein lactylation via intracellular metabolism and the glycolytic pathway. To test this hypothesis *in vivo*, we treated mice with electroconvulsive stimulation (ECS) and then examined lactate and protein lactylation levels in the brain. The brain lactate levels increased 10 minutes after ECS and persisted for 1 hour (Fig. 2d, e). We then examined Kla levels in mouse brains treated with ECS in combination with OX and/or 4-CIN (Fig. 2f). While the ECS-induced increase in Kla immunoreactivity was not significantly affected by prior treatment with either OX or 4-CIN alone, it was attenuated by a combination treatment with OX and 4-CIN (Fig. 2g, h), suggesting that, *in vivo*, both intracellular new lactate production and extracellular lactate uptake have a role in neuronal activation-induced protein lactylation in the acute phase.

**Figure 2.**
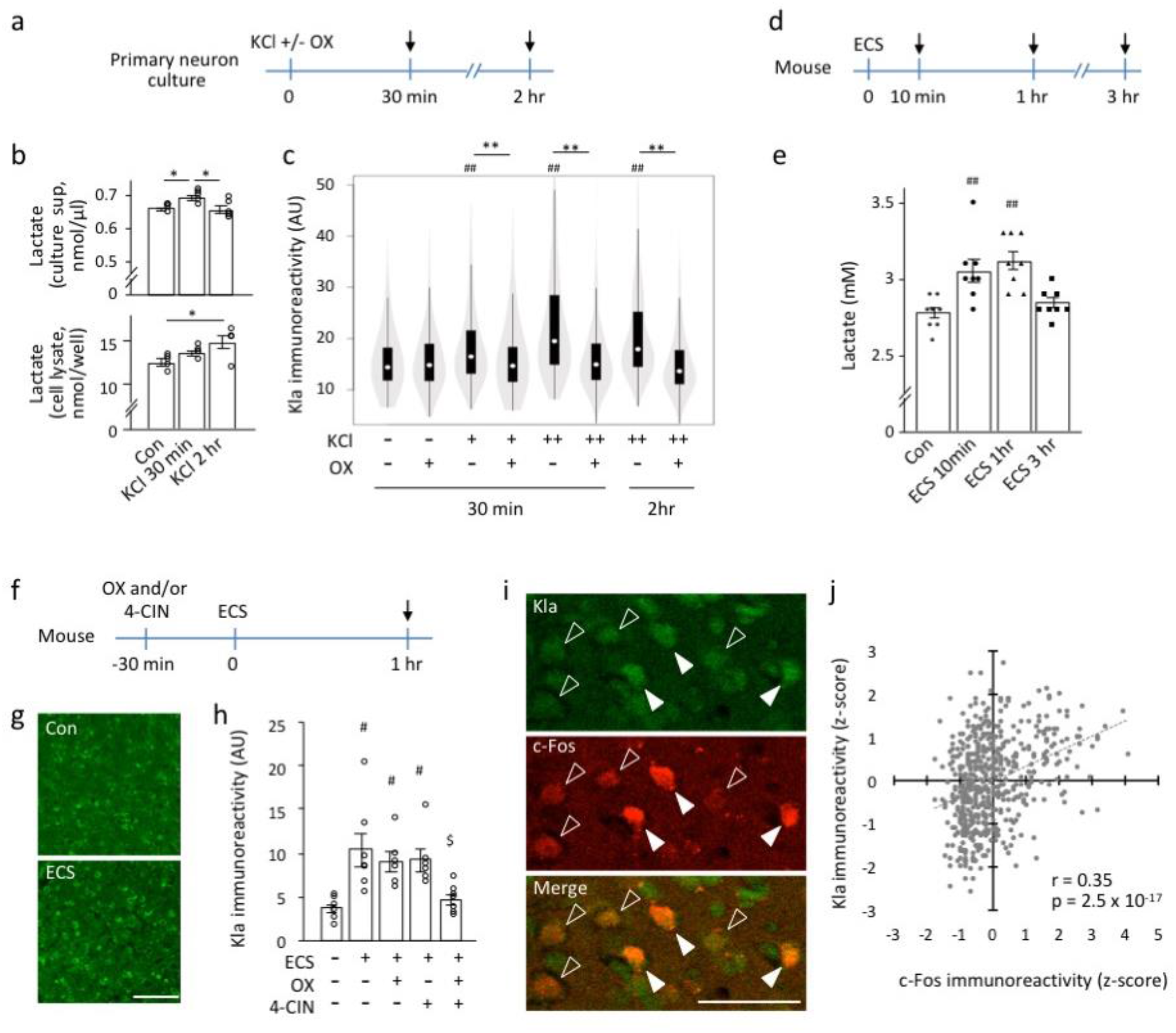
Neuronal excitation stimulates protein lactylation in neurons. (a–c) Primary cultured cortical neurons (10 DIV) were treated with potassium chloride (KCl), with or without sodium oxamate (OX). Arrows indicate sampling time points (a). Cultured neurons were treated with KCl (100 mM) and lactate concentrations were measured in the culture supernatant (upper panel, b) and cell lysate (lower panel, b). *P < 0.05, one-way analysis of variance (ANOVA) followed by post-hoc pairwise comparisons. N = 5 for each condition. Statistical results are provided in the Extended Data Table 5. (c) Results of Kla immunostaining analysis of cultured neurons expressing βIII-tubulin treated with KCl [10 mM (+) or 100 mM (++)] and OX [20 mM (+)] (c) At least 300 cells were examined for each condition. **P < 0.01; ##P < 0.01 compared to the KCl/OX condition; one-way ANOVA followed by post-hoc pairwise comparisons (Extended Data Table 6). (d, e) Brain lactate levels in mice treated with electroconvulsive stimulation (ECS) (N = 8 mice for each condition). ##P < 0.01 compared to control; one-way ANOVA followed by post-hoc pairwise comparisons (Extended Data Table 7). (f–h) Kla in the mouse brain treated with ECS. Adult mice were treated with ECS 30 minutes after treatment with OX, α-cyano-4-hydroxycinnamate (4-CIN), or saline. Brain sampling was performed 1 hour after ECS treatment (arrow, f). Immunostaining images of Kla in the prefrontal cortex (PFC). Scale bar: 100 μm (g). Graph showing Kla immunoreactivity in the PFCs of the mice (h). #P < 0.05 compared to ECS-/OX-/4-CIN-, $P < 0.05 compared to ECS+/OX-/4-CIN-; one-way ANOVA followed by post-hoc pairwise comparisons. N = 6–7 mice for each condition (Extended Data Table 8). (i) Kla and c-Fos double immunostaining images of the mouse PFC at 1 hour after an ECS. Open and solid arrowheads indicate cells with low and high levels of Kla + c-Fos, respectively. Scale bar: 50 μm. (j) Scatterplot showing a correlation between Kla and c-Fos immunoreactivity in individual cells in the PFC (represented as Z-score transformed values). Data from 543 cells from 15 mice in four different conditions (Extended Data Fig. 4) are integrated in the graph.

c-Fos is an immediate early gene product used as a marker for neuronal activation^15^. Utilizing the characteristics of c-Fos expression, we investigated the relation between Kla and c-Fos expressions in the ECS mouse model. Upon double-immunostaining analysis of c-Fos-expressing cells in the PFC, we found a significant positive correlation between Kla and c-Fos immunoreactive intensities on an individual cell basis level (Fig. 2i, j, Extended Data Fig. 4). Considering that intensity of electrical stimulation positively correlates with c-Fos-labeling levels in individual neurons^16^, our results suggest that the more neurons are activated, the more Kla is induced in the cells. However, we have to note that there also are many cells that are Kla-positive but c-Fos-negative, suggesting that some Kla formations are independent of neuronal activation or lasts longer than that of c-Fos protein expression after neuronal activation.

### Social stress stimulates protein lactylation in the brain

We explored whether protein lactylation in the brain is modified by a physiologically relevant stimulus using a social defeat stress (SDS) model of depression. SDS has been shown to increase c-Fos expression levels in many brain regions^17–19^. Brain lactate levels increased immediately (10 minutes) after a single exposure to the stress, and the increase was attenuated by prior treatment with OX^1^ (Extended Data Fig. 5).

Further, we conducted a chronic experiment where the mice experienced daily defeat stress with prior treatment with OX or vehicle (saline) for 10 consecutive days. To examine the correlation between brain lactate and Kla levels and behavioral measures of anxiety and depression, we conducted a series of behavioral tests prior to brain sampling (Fig. 3a, Extended Data Fig. 6, 7). In the social avoidance test, the social interaction ratio decreased in both chronic vehicle-treated and OX-treated socially defeated mice compared to control mice (Fig. 3b). Based on the standard criteria^20,21^, we focused on mice susceptible to SDS, which showed a social interaction ratio <1 in the social avoidance test and designated them as defeated mice for the following behavioral and brain tissue analyses.

**Figure 3.**
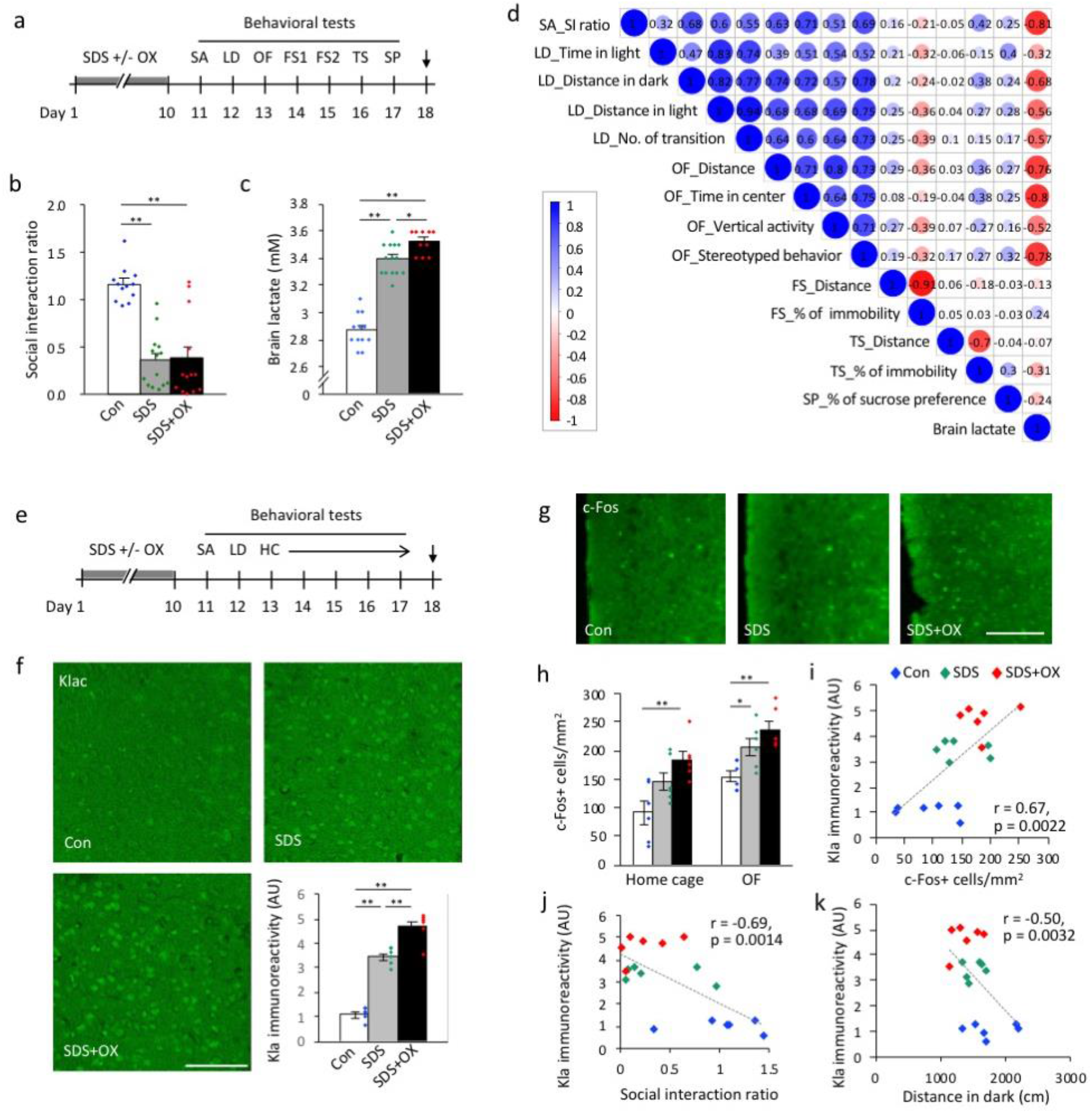
Social stress induces protein lactylation in the brain. (a–c) Mice were exposed to repeated social defeat stress, with or without sodium oxamate injections and then analyzed by a series of behavioral tests (a). Arrow in (a) indicates the sampling time point. FS: Porsolt forced swim test; LD: light-dark transition test; OF: open field test; OX: sodium oxamate; SA: social avoidance test; SDS: social defeat stress; SP: sucrose preference test; TS: tail suspension test. (b) Social avoidance test. (c) Brain lactate levels in the three groups of mice. N = 10–14 mice for each condition. Statistical results are provided in the Extended Data Table 9 and 10, respectively. (d) Correlation matrix showing correlation coefficients between behavioral measures and brain lactate levels in control and socially defeated mice. (e, f) Brain sampling for immunohistochemical analysis was performed using another set of mice exposed to SDS with or without OX injection. (e) HC: home cage locomotor activity monitoring test. (f) Immunostaining images of Kla and bar graph showing Kla immunoreactivity in the prefrontal cortex (PFC). Scale bar: 100 μm. **P < 0.01, one-way analysis of variance followed by post-hoc pairwise comparisons; N = 6 mice for each condition (Extended Data Table 11). (g) Immunostaining images of c-Fos in the PFC. (h) Bar graphs showing c-Fos-positive cells in the PFC (Extended Data Table 12). Mice in the home cage group were sampled immediately after being removed from the home cage. Mice in the OF group were sampled after a 2-hour exposure to the open field. (i–k) Scatterplots showing correlations between Kla immunoreactivity and c-Fos expression in the PFC and (i) the social interaction ratio in the social avoidance test (j), and the distance traveled in the dark compartment in the light/dark transition test, in mice from the OF group. N = 6 for each condition.

We found that brain lactate levels were higher in defeated mice than in control mice (Fig. 3c), as confirmed by independent sets of mice (Extended Data Fig. 8). Brain lactate levels were paradoxically increased in OX-treated defeated mice at 9 days after the last stress exposure and OX treatment compared to vehicle-treated defeated mice and control mice (Fig. 3c). Simple correlation (Fig. 3d; Extended Data Fig. 9) and multiple regression (Extended Data Table 1) analyses suggested that increased brain lactate levels are preferentially associated with increased anxiety-like behaviors. In this stress model, Kla levels in the PFC were increased in vehicle-treated defeated mice and, to a larger extent, in OX-treated defeated mice compared to control mice (Fig. 3e, f).

Expression of neuronal activity marker c-Fos in the PFC also increased in vehicle-treated defeated mice and, to a lager extent, in OX-treated defeated mice, in both a familiar environment in the home cage (non-stimulated basal condition) and novel environment exposed to the open field (condition with anxiogenic stimulation) (Fig. 3g, h). In the mice that received anxiogenic stimulation to ensure more c-Fos-expressing cells, we found that the Kla level positively correlated with the c-Fos expression level in the PFC (Fig. 3i). Although changes in Kla and c-Fos expression levels were observed in some other brain regions implicated in anxiety and depression behaviors (e.g., hippocampus, lateral habenula, and amygdala; Extended Data Fig. 10, 11), no significant correlations were found between Kla and c-Fos expression levels in brain regions other than the PFC (Extended Data Fig. 12), suggesting that simultaneous induction of c-Fos and Kla occurs in a brain region-specific manner following repeated SDS. Furthermore, we found that Kla levels in the PFC, but not in the amygdala and hippocampal subregions, negatively correlated with the social interaction ratio in the social avoidance test (Fig. 3j, Extended Data Fig. 13) and with distance traveled in the dark compartment in the light–dark transition test (Fig. 3k), indicating that the more susceptible to SDS and the more anxious the mouse is, the higher the Kla levels in the PFC.

### Identification of lactylated proteins in the brain

We sought to identify and quantify the brain proteins modified by lactylation and whose lactylation levels changed in response to SDS. Accordingly, we analyzed the PFC of defeated and control mice using Kla antibody immunoprecipitation and mass spectrometry techniques. This analysis identified 63 candidate lysine-lactylated proteins in the mouse PFC (Extended Data Table 2). GO-enrichment analysis with DAVID^22,23^ showed that these proteins were mainly enriched in nucleosome- and histone-related biological processes (Extended Data Table 3). Among them, 12 proteins were suggested to be changed by SDS, of which seven were upregulated and five were downregulated. Notably, four subtypes of histone H1 were included in the 12 changed proteins and all were upregulated, suggesting that SDS increases lactylation at histone H1. We confirmed these observations by conducting immunoprecipitation with the histone H1 antibody followed by western blot analysis with the Kla antibody (Extended Data Fig. 14). In the PFC, 59.5% of cells were positive for histone H1 and nearly all the histone H1-positive cells (94.6%) were Kla positive, and vice versa (99.1% of Kla-positive cells were histone H1 positive) (Fig. 4a, b). Upon super-resolution imaging using stimulated emission depletion (STED) microscopy, we observed histone H1 and Kla double-positive structures in the nucleus, suggestive of lysine lactylation at histone H1 (Fig. 4c). Importantly, the double-positive structures in the nucleus increased in the defeated mice compared to control mice (Fig. 4d), further supporting that SDS enhances lysine lactylation levels at histone H1 in the PFC.

**Figure 4.**
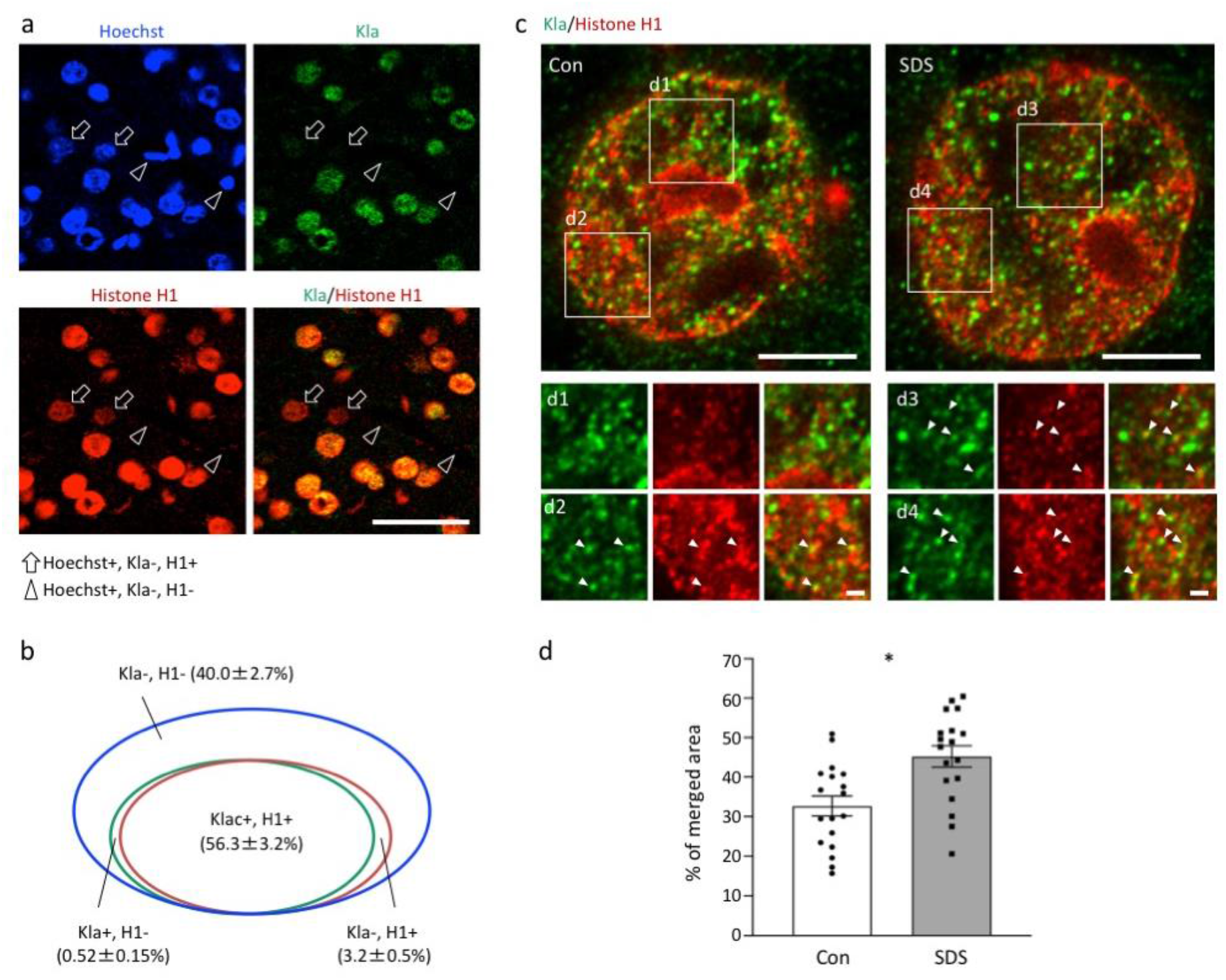
Histone H1 lactylation in the brain is increased by social stress. (a) Double immunostaining of Kla and histone H1 in the mouse PFC. Arrows indicate Kla-negative and histone H1-positive Hoechst-stained cells, and arrowheads indicate Kla-negative and histone H1-negative Hoechst-stained cells. Scale bar: 50 μm. (b) Venn diagram showing the overlap between Kla-positive and histone H1-positive cells in the mouse PFC. (c) Super-resolution images of Kla and histone H1 in the nuclei of neurons of socially defeated mice recorded with STED microscopy. Arrowheads indicate Kla- and histone H1-double–positive structures. Scale bars: 5 μm (upper panels), 1 μm (lower panels). (d) Bar graphs show the percentage of Kla and histone H1 merged area in the nucleus. N = 18 cells from 3 mice for each condition. *P = 0.012, linear regression analysis.

## Discussion

In this study, we showed that a novel post-translational modification called lysine lactylation occurs in brain cells and that its levels can be changed by neuronal excitation. We found that increasing lactate levels by lactate addition or inducing neuronal excitation with high-K or ECS increased Kla levels in the mouse brain cells both *in vitro* and *in vivo*. The lactate and Kla levels also increased by behaviorally induced neuronal excitation associated with SDS. In the SDS model of depression, Kla levels in the PFC, a core brain region implicated in depressive and anxiety disorders, correlated with anxiety-like behaviors, suggesting a potential behavioral significance of Kla in the brain. Furthermore, our proteomic approach identified 63 candidate lactylated proteins in the mouse PFC and highlighted that histone H1 lactylation increases in response to chronic stress.

Our results using SDS model mice suggest that, in stress-related pathological conditions, excess neuronal activity may underlie increased protein lactylation in the PFC, potentially associated with susceptibility to stress and anxiety-like behavior, although the causal mechanism remained unclear. Given that increased Kla levels reflect neuronal activity, it might be expected that significant correlations can be seen between Kla levels in the brain regions other than PFC and behavioral alterations. Adaptation in neuronal activity has been suggested to occur following repeated SDS in some brain regions; for example, neurons in the central nucleus of amygdala became less active and showed suppressed c-Fos expression after repeated SDS^24^. Such adaptation in neuronal activity may affect c-Fos expression, possibly as well as Kla induction, resulting in no clear correlations between behavior and Kla levels in brain regions other than the PFC.

Regarding the mechanisms of Kla formation, Zhang et al. showed that, using a ^13^C-labeled L-lactate incorporation method, Kla could be derived directly from lactate exogenously applied to mouse bone marrow-derived macrophages^1^. Conversely, other studies have suggested that Kla formation could be mediated indirectly by a glycolytic intermediate, lactoylglutathione, which can provide a lactyl group to lysine residues. L-Lactate is suggested to suppress the activity of glyoxalase 2, an enzyme catalyzing the conversion of lactoylglutathione into glutathione and free D-lactate. Such suppression due to increased L-lactate could elevate lactoylglutathione levels, which in turn, may facilitate the indirect lactylation pathway. Gaffney et al. used a stable isotope labeling of amino acids in cell culture (SILAC) model to identify lactylated proteins that could be formed via lactoylglutathione in the HEK293 cell line^25^. The identified proteins were enriched in glycolytic and carbon-metabolism enzymes but did not include histone family proteins. As mentioned above, Kla forms directly from lactate on histone H3^1^. Our list includes histone H3 as well as the 60S ribosomal protein L30 and Rho guanine nucleotide exchange factor 2, also found in the 350-protein list published by Gaffney et al.^25^ These observations suggest that direct and indirect lactylation pathways may coexist, rather than being exclusive, depending on the proteins expressed in the brain. Regardless of the mechanisms, lysine lactylation may play a role in coupling metabolic changes to gene-expression regulation.

Our analyses highlighted that H1 histones increase their Kla levels in response to SDS. Histone H1 is known to be a linker histone, having a key role in higher-order chromatin folding and genome stability. Seven subtypes of H1 histone have been identified in mammalian somatic cells, i.e., H1.0–1.5 and H1.10^26^. Numerous post-translational modifications have been identified in H1, such as phosphorylation, methylation, acetylation, formylation, crotonylation, and citrullination^27^. Post-translational modifications of histone H1 have received less attention than those of core histones; therefore, knowledge on the functional role of H1 modifications is limited^26,28^, especially in brain cells. In the case of mouse macrophage histone H3, lactylation competes with acetylation at lysine residues to regulate the expression of a specific gene set. Extrapolating these findings to histone H1 and considering that histone H1 acetylation regulates chromatin compaction and decondensation and hence the accessibility of linker DNA regions to transcription factors, histone H1 lactylation may also be involved in gene-expression regulation depending on the balance of Kla with acetylation.

In conclusion, this study indicates that a novel post-translational modification called lysine lactylation occurs in the brain, and it is regulated by neuronal activity. While 63 candidate lactylated proteins were identified in the brain, it remains unclear whether such a modification has functional significance for each protein at the cellular and behavior levels, and if it does, in what brain regions and in what cell types and in what types of behavioral phenotypes. It would also be interesting to explore whether protein lactylation in the brain is related to the etiology and pathophysiology of neuropsychiatric disorders, as are altered lactate levels in the brains of animals^3^. Further studies may reveal unknown roles of lactate, linking neural activity, glycolytic metabolism, and molecular signaling mediated by protein lactylation under physiological and pathophysiological conditions.

## Methods

### Animals

C57BL/6J mice (7–9 weeks old) were purchased from Charles River Laboratories Japan, Inc. (Hino, Japan) and Japan SLC (Shizuoka, Japan). Retired ICR mice were purchased from Chubu Kagaku Shizai Co. Ltd. (Nagoya, Japan) and Japan SLC. The mice were habituated to their environments for a week before experimentation. All animal experiments were approved by the Institutional Animal Care and Use Committee of Fujita Health University, Ibaraki University, and The National Institute of Advanced Industrial Science and Technology based on the Law for the Humane Treatment and Management of Animals and the Standards Relating to the Care and Management of Laboratory Animals and Relief of Pain. Every effort was made to minimize the number of animals used.

### Cell culture

Embryonic C57BL/6J mice (15.5–16.5 days old) were used for primary neuron culture. The frontal cortices and hippocampi were dissected from the brains in cold HEPES-buffered, calcium and magnesium-free Hank’s balanced salt solution (HBSS; 14175-103, Thermo Fisher Scientific, Waltham, MA, USA) and dissociated in HBSS containing 0.25% papain (LS003127, Worthington Biochemical Corporation, Lakewood, NJ, USA), 0.1% L-cysteine (C7352, Merck, Darmstadt, Germany), 0.1% bovine serum albumin (BSA; A2153, Merck), and 0.01% DNase I (D4263, Merck) at 37°C for 12 minutes. The enzymatic digestion was stopped by adding a plating medium (Dulbecco’s modified Eagle medium, D5796, Merck, containing 10% fetal bovine serum). Tissues were dissociated by pipetting, centrifuged at room temperature for 10 minutes, and resuspended in the plating medium. Cells were counted, seeded at a density of 25,000 cells/cm^2^ on culture plates or glass coverslips previously coated with 0.01% poly-L-lysine (P8920, Merck), and incubated at 37°C. After 2 hours, the plating medium was replaced by a culture medium (Neurobasal™ medium [21103-049, Thermo Fisher Scientific] containing B-27™ Supplement [17504-044, Thermo Fisher Scientific], 0.5 mM L-glutamine [G7513, Merck], and 1% penicillin-streptomycin solution [P4333, Merck]). The culture medium was changed by one half with fresh medium every 3–4 days. At 10–14 days *in vitro*, drug treatment experiments, immunocytochemical analysis were conducted.

### Immunocytochemistry

Cells were fixed with 4% paraformaldehyde (PFA) in phosphate-buffered saline (PBS) for 10 minutes, pre-incubated for 30 minutes in 5% BSA (Merck) in PBS containing 0.05% Tween-20 (PBST), and then incubated for 30 minutes in PBS containing the primary antibodies: rabbit polyclonal anti-lactyllysine antibody (PTM-1401, PTM Biolabs, Hangzhou, China), mouse monoclonal anti-tubulin β3 (801201, BioLegend, San Diego, CA, USA), and mouse monoclonal anti-glial fibrillary acidic protein antibody (G3893, Sigma-Aldrich, St Louis, MO, USA). Immunoreactivity to the antigen was visualized using Alexa Fluor 488- and Alexa Fluor 594-conjugated secondary antibodies (Molecular Probes, Eugene, OR). Nuclear staining was performed with Hoechst 33258 (Polysciences, Warrington, PA). All reactions were performed at room temperature. We used a microscope (LSM 510 META; Zeiss, Göttingen, Germany) to obtain images of the stained cells. The soma of tubulin β3-positive cells was manually delineated and the signal intensity of Kla staining within the region of interest was recorded using ZEN software (Zeiss). At least 200 cells from triplicate wells per condition were analyzed.

The specificity of the Kla antibody was confirmed using dot blot analysis of lysine residues with various types of modifying groups (i.e., acetyl-, crotonyl-, propionyl-, butyryl-, trimethyl-, and succinyl-lysine); the antibody showed almost no cross-reactivity with these modified lysine residues^1,29^. Additionally, lactate treatment dramatically increased Kla immunoreactivity in human MCF-7, Hela, and MDA-MB-231 cell lines^1,29^.

### Immunohistochemistry

Immunohistochemical analysis was performed as previously described with some modifications^30,31^. Briefly, mice were deeply anesthetized with isoflurane and transcardially perfused with PBS followed by 4% PFA in PBS. The brains were dissected, immersed overnight in the same fixative, and cryoprotected by sequential incubation in 10, 20, and 30% sucrose in PBS for 2–3 days each at 4°C. After cryoprotection, brains were mounted in Tissue-Tek optimal cutting temperature compound (Miles, Elkhart, IN), frozen, and cut into 30-μm-thick coronal sections using a microtome (CM1850; Leica Microsystems, Wetzlar, Germany). The sections were stored in PBS containing 0.1% sodium azide at 4°C until further use. The sections were pre-incubated for 1 hour at room temperature in 5% skim milk in PBST and then incubated overnight at 4°C in PBS containing the primary antibodies. We used rabbit polyclonal anti-lactyllysine antibody (PTM-1401, PTM Biolabs), mouse monoclonal anti-CaM Kinase II α subunit antibody (05-532, Merck), mouse monoclonal anti-GAD67 antibody (MAB5406, Millipore), mouse monoclonal anti-S-100 β subunit antibody (S2532, Merck), rat monoclonal anti-CD68 antibody (ab53444, abcam), and mouse monoclonal anti-c-Fos antibody (sc-166940, Santa Cruz Biotechnology, Santa Cruz, CA, USA). For c-Fos staining, antigen retrieval was performed by autoclave heating at 120°C for 1 minute in 0.01 M sodium citrate buffer prior to the blocking step. Immunoreactivity to the antigen was visualized using Alexa Fluor 488-or Alexa Fluor 594-conjugated secondary antibody (Molecular Probes). Nuclear staining was performed with Hoechst 33258 (Polysciences). We used a microscope (LSM 510 META; Zeiss) to obtain images of the stained sections. Three to seven sections from each animal were processed for semi-quantification analyses in each brain region, and the averaged values were considered as one sample. c-Fos-positive cells were counted in the indicated brain regions, manually delineated using ZEN software (Zeiss) according to the mouse brain atlas^32^. For c-Fos and Kla immunofluorescence intensity analysis in individual cells, c-Fos-positive cells were circled, and the signal intensity for each antibody within the circled areas was recorded using Zen software (Zeiss). For Kla immunofluorescence intensity, to reduce potential artifacts across sections, the values were calculated from the mean signal intensity of the entire image by subtracting background signals using Zen software (Zeiss). Semi-quantification analyses were performed in the medial PFC (approximately from 2.1 to 2.4 mm from the bregma), dorsal hippocampus, lateral habenula, and amygdala (approximately from −2.1 to −1.6 mm from the bregma), and ventral hippocampus (approximately from −3.5 to −3.8 mm from the bregma).

### STED microscopy

Brain slices were doubleimmunostained as described above. Mouse monoclonal anti-histone H1 antibody (sc-8030; Santa Cruz Biotechnology, Inc.) and rabbit polyclonal anti-Kla antibody were the primary antibodies, and Alexa Fluor 555- and Alexa Fluor 488 conjugated antibodies (Molecular Probes) were the secondary antibodies.

STED images were acquired with a TCS SP8 STED 3X system (Leica Microsystems, Wetzlar, Germany) with 2ch HyDSMD detectors^33^. A white laser (for excitation) and a 660-nm laser with a donut beam (for stimulated emission depletion) were used for STED imaging. For detection of histone H1 and Kla, we used excitation wavelengths of 561 and 488 nm, respectively. Raw STED images (pixel size: 20 nm) were recorded with a 93× glycerol immersion objective (Leica HC PL APO 93×/1.30 GLYC motC STED W) and deconvoluted using the Huygens software (SVI, Hilversum, Netherlands). The percentage of merged area in a nucleus stained for both histone H1 and Kla was estimated using the ImageJ program.

### Immunoprecipitation

The PFCs were dissected, snap-frozen in liquid nitrogen, and stored at −80°C until further use. Fifty microliters of Dynabeads™ Protein G solution (10007D; Thermo Fisher Scientific) was separated into low-protein absorption tubes (PROKEEP; Watson Bio Lab, Tokyo, Japan) and reacted with 2 μg of rabbit polyclonal anti-lactyllysine antibody (PTM Biolabs) or normal rabbit IgG (Cell Signaling Technology, Inc., Danvers, MA, USA). The frozen tissues were then homogenized in lysis buffer (FNN0021, Thermo Fisher Scientific). Then, 200 μL of 1-μg/μL tissue lysate was added to the bead-antibody complex and incubated with a rotation of 30 minutes at room temperature. The bead-antibody-antigen complex was then processed for mass spectrometry.

### Mass spectrometry

#### Trypsin digestion

Bead-antibody-antigen complex was washed with 50 mM ammonium bicarbonate. The binding proteins were eluted by trypsin (Promega Corporation, Madison, WI, USA) for 1 hour at 37°C, reduced with 10 mM dithiothreitol for 30 minutes at room temperature, and alkylated with 20 mM iodoacetamide for 30 minutes at room temperature, followed by an overnight incubation at 37°C. Digestion was stopped by adding trifluoroacetic acid. The digested sample was desalted using GL-Tip SDB (GL Sciences, Tokyo, Japan) and then dissolved in 20 µL of 0.1% trifluoroacetic acid.

#### Liquid chromatograph-tandem mass spectrometer (LC-MS/MS) analysis

The peptides were analyzed using LC-MS (EASY-nLC™ 1000, Orbitrap Fusion EDT; Thermo Fisher Scientific). Injected peptides were trapped on an Acclaim™ PepMap™ 100 C18 LC column (3 µm particle size, 0.075 mm inner diameter × 20 mm length; Thermo Fisher Scientific) and separated on an EASY-Spray LC column (3 µm particle size, 0.075 mm inner diameter × 150 mm length; Thermo Fisher Scientific) with a linear gradient of 0–35% acetonitrile with 0.1% formic acid over 60 minutes at a flow rate of 300 nL/minute. Data acquisition was performed using Xcalibur™ (Thermo Fisher Scientific). Full MS spectra (m/z 375–1500) were acquired at a resolution of 120,000. The most intense precursor ions were selected for collision-induced fragmentation in the linear ion trap at a normalized collision energy of 35%. Dynamic exclusion was employed within 60 seconds to prevent repetitive selection of peptides.

#### Data analysis

Raw data were analyzed using Proteome Discoverer™ (version 2.2, Thermo Fisher Scientific) with the Mascot search engine (version 2.6.1, Matrix Science Ltd., London, UK) against the Swiss-Prot database. The search parameters were as follows: mass tolerances in MS and MS/MS, 10 ppm and 0.6 Da; maximum missed cleavages allowed, 2. Methionine oxidation and N-acetylation were set as variable modifications, and cysteine carbamidomethylation was set as a fixed modification. A target-decoy database search was used to estimate the false-positive ratio for peptide identification; the false discovery rate was set at 1%.

### Western blot analysis

Western blot analysis was performed as previously described, with some modifications^30,31^. Tissues from the brain and peripheral organs were collected from adult c57BL6/J mice, snap-frozen in liquid nitrogen, and stored at −80°C until further use. Peritoneal cavity cells were collected from adult C67BL/6J mice and used as a macrophage sample. Tissues and cultured cells were homogenized in lysis buffer (MCL1, Merck). Protein homogenates were separated with gel electrophoresis using NuPAGE™ 4–12% Bis-Tris gels (Thermo Fisher Scientific) and transferred to polyvinylidene difluoride membranes. Membranes were preincubated in 5% skim milk in Tris-buffered saline with 0.05% Tween® 20 (TBST) for 1 hour and incubated for 2 hours in primary antibody, rabbit polyclonal anti-lactyllysine antibody (PTM Biolabs), or mouse monoclonal anti-β-actin antibody (A5316, Merck) diluted in TBST. They were then incubated for 1 hour with a secondary antibody conjugated with horseradish peroxidase (sc-2357, anti-rabbit IgG; sc-516102, anti-mouse IgG; Santa Cruz Biotechnology) diluted with TBST. All reactions were performed at room temperature. Immunoreactivity was visualized with a chemiluminescence detection regent (ImmunoStar LD, 292-69903; FUJIFILM Wako Pure Chemical Corporation, Osaka, Japan) and photographed with a luminescent image analyzer (LAS-4000 mini; GE Healthcare, Buckinghamshire, UK).

### Electroconvulsive stimulation

The experimental mice were briefly anesthetized with isoflurane, and bilateral electroconvulsive stimulation (ECS; current: 25 mA; shock duration: 1 s; frequency: 100 pulses/sec; pulse width: 0.5 msec) was administered via ear clip electrodes with a pulse generator (ECT Unit; Ugo Basile, Gemonio, VA, Italy)^34^. Control mice were handled similarly, except that they were not administered ECS.

### Chronic SDS

#### Condition 1

SDS serves as a preclinical model of chronic stress^35,36^. We performed chronic social defeat stress in mice at Fujita Health University, as previously described^36^.

##### Screening of aggressor ICR mice

Retired ICR breeders were screened for aggressive behavior for 3 consecutive days. They were individually housed in a clear rectangular hamster cage (26.7 cm [width] × 48.3 cm [depth] × 15.2 cm [height]; Allentown Caging Equipment, Allentown, NJ, USA) equipped with a clear perforated Plexiglass divider (0.6 cm [width] × 45.7 cm [depth] × 15.2 cm [height]; Chubu Kagaku Shizai [custom order]) and a steel-wire top (Allentown Caging Equipment).

##### Chronic SDS

Experimental C57BL/6J mice were placed in the compartment containing an aggressor ICR mouse and were exposed to social defeat for 10 minutes. During this period, the experimental mice generally displayed submissive behaviors. They were then transferred to the other compartment of the aggressor’s cage, allowing sensory interaction with the intruder C57BL/6J mouse and resident ICR aggressor mouse for approximately 24 hours (cohabitation housing condition). This procedure was repeated for 10 consecutive days, with each experimental C57BL/6J mouse being exposed to a novel ICR aggressor each day to prevent habituation. Control C57BL/6J mice were placed in pairs in an identical home cage separated by a perforated divider. The control animals were rotated to a new cage compartment daily and allowed to interact with a novel mouse of the same strain. Following the last defeat session, all C57BL/6J mice were singly housed in standard mouse cages. Twenty-four hours after the last defeat session, a social avoidance test and other behavioral tests were conducted as described. Brain sampling was performed 8 days after the last defeat session. *Condition 1* was used for OX-treatment experiments.

#### Condition 2

The experiment was performed at Fujita Health University, as described in *Condition 1*, except that the intruder C57BL/6J mice were individually housed in separate cages, away from ICR aggressor mice, after the 10-minute-aggression (separation housing condition). As in *Condition 1*, brain sampling was performed 8 days after the last defeat session.

#### Condition 3

The experiment was performed at Ibaraki University, as previously described^37^. The intruder C57BL/6J mice were exposed to sensory interaction with a resident ICR aggressor mouse in the other compartment of the aggressor’s cage for approximately 24 hours (cohabitation housing condition). Twenty-four hours after the last defeat session, a social avoidance test was conducted ^37^. Brain sampling was performed 2 days after the last defeat session.

### Drug treatment

Twenty minutes prior to each defeat session, the mice were intraperitoneally administered sodium oxamate (OX; 1 g/kg; O2751, Sigma-Aldrich, Tokyo, Japan) or sodium L-lactate (1 g/kg; L-7022, Sigma-Aldrich). They were administered the same amount of saline as the control. OX is an inhibitor of lactate dehydrogenase (LDH) activity in the central nervous system^13,14^. The dose of OX that we used was previously shown to be effective in suppressing seizures in a mouse model of epilepsy^14^. The dose of sodium L-lactate that we used was previously shown to exert antidepressant-like effects in a corticosterone-induced chronic stress model in mice^7^.

### Behavioral tests

Most of the behavioral tests were performed as described elsewhere^38–41^, unless otherwise noted.

#### Social interaction test

Social interaction was tested 1 day after the last defeat session as previously described^36^, with minor modifications. We used an open field chamber (40 cm (width) × 40 cm (depth) × 30 cm (height); O’HARA & Co., Tokyo, Japan) and a wire mesh cage (10.5 cm (width) × 6.5 cm (depth) × 29 cm (height); O’HARA & Co.) to enclose the ICR aggressor mouse. The ICR aggressor mouse that is novel to the defeated C57BL/6J mouse was used. Each social interaction test was composed of two 150-second phases: the ICR aggressor mouse is absent and present in the interaction zone for the first and second phase, respectively. An identical, empty wire mesh cage was placed for the first phase testing. A 13.7 cm × 23.6 cm rectangular zone including the wire mesh cage was defined as the social interaction zone. Mouse behaviors were video-monitored, and the trajectory of mouse ambulation, distance traveled, and time spent in the interaction zone for all test phases were automatically determined using ImageSI (see “Image analysis”). A social interaction ratio was calculated by dividing the time spent in the interaction zone when the target was present by the time spent in the interaction zone when the target was absent. A social interaction ratio of 1 was considered the threshold for dividing defeated mice into the susceptible and resilient groups.

#### Light/dark transition test

A light/dark transition test was conducted as previously described^38–42^. The apparatus consisted of a cage (42 cm (width) × 21 cm (depth) × 25 cm (height)) divided into two sections of equal size by a partition with a door (O’HARA & Co.). One chamber was brightly illuminated (390 lux), whereas the other chamber was dark (2 lux). Mice were placed into the dark chamber and allowed to move freely between the two chambers with the door open for 10 minutes. The total number of transitions, latency to first enter the lit chamber, distance traveled, and time spent in each chamber were recorded using ImageLD (see “Image analysis”).

#### Open field test

The apparatus was a transparent square cage (42 cm (width) × 42 cm (depth) × 30 cm (height)) equipped with infrared photobeam sensors (VersaMax; Accuscan Instruments, Columbus, OH, USA) with a white floor^38–41^. The center of the floor was illuminated at 100 lux. Each mouse was placed in the corner of the cage. The total distance traveled (cm), vertical activity (rearing measured by counting the number of photobeam interruptions), time spent in the center area (20 cm × 20 cm), and stereotypic counts (defined by the number of breaks of the same beam) were recorded for 30 minutes using the VersaMax system.

#### Porsolt forced-swim test

A Plexiglas cylinder (20 cm height × 11.4 cm diameter; O’HARA & Co.) filled with water (21–23 °C) up to a height of 8 cm was placed in a white plastic chamber (32 cm (width) × 44 cm (depth) × 49 cm (height); O’HARA & Co.) ^38–41^. Mice were placed into the cylinder, and both immobility and distance traveled were recorded over a 10-minute test period. The test was conducted for 2 consecutive two days, and the results of the first trial are shown. Images were captured at two frames per second. For each pair of successive frames, the amount of area (pixels) in which the mouse moved was measured. When the amount of area was below a certain threshold, mouse behavior was classified as “immobile.” When the amount of area equaled or exceeded the threshold, the mouse was classified as “moving”. The optimal threshold to judge movment was determined by adjusting it to the amount of immobility measured by human observation. Immobility lasting for less than 2 seconds was not included in the analysis. Data acquisition and analysis were performed automatically, using ImagePS/TS (see “Image analysis”).

#### Tail suspension test

Each mouse was suspended 30 cm above the floor by the tail in a white plastic chamber (41cm (width) × 31 cm (depth) × 41 cm (height); O’Hara & Co.)^38–41^. The behavior was recorded for 10 minutes. Images were captured through a video camera, and immobility was measured. Similar to the Porsolt forced swim test mentioned above, immobility was judged using ImagePS/TS (see “Image analysis”) according to a certain threshold. Immobility lasting <2 seconds was not included in the analysis.

#### Sucrose preference test

C57BL/6J mice were subjected to a two-bottle choice test, in which mice were provided with a bottle containing water and a second bottle containing 1% sucrose solution, with the left/right positions of the bottles counterbalanced across groups ^39,40^. The mice were habituated to a two-bottle condition during all the experimental period. Sucrose preference during 4-hour in the test day was calculated as the percentage of sucrose preference = 100 × {[sucrose intake (g)]/(sucrose intake (g) + water intake (g)]).

#### Locomotor activity monitoring in the home cage

Locomotor activity monitoring in the home cage was performed as previously described^30,41^. A system that automatically analyzes the locomotor activity of mice in their home cage was used^43^. The system contains a home cage (29 cm (width) × 18 cm (depth) × 12 cm (height)) and filtered cage top, separated by a 13-cm-high metal stand containing an infrared video camera, attached to the top of the stand. Each mouse was individually housed in each home cage, and their locomotor activity was monitored for 6 days. Mice were allowed to habituate to the test apparatus, and hence, the data of the latter 3 days were analyzed. Distance traveled was measured automatically using ImageHA (see “Image analysis”).

#### Image analysis

For the social interaction test, light/dark transition test, Porsolt forced swim test, and tail suspension test, image analysis programs (ImageSI/LD/PS/TS/HA) were used to automatically analyze mouse behaviors. The application programs, based on the public domain ImageJ program (developed by Wayne Rasband at the National Institute of Mental Health, Bethesda), were developed and modified for each test by Tsuyoshi Miyakawa. ImageLD program can be freely downloaded from the website of the “Mouse Phenotype Database” (http://www.mouse-phenotype.org/)^44^.

### Measurements of brain lactate levels and pH

In the *in vivo* experiments, the whole brain was used to measure tissue pH and lactate levels as previously described^3^. Briefly, snap-frozen tissues were homogenized in ice-cold distilled H_2_O (5 mL per 500 mg of tissue). The pH of the homogenates was measured using a pH meter (LAQUA F-72, HORIBA, Ltd., Kyoto, Japan) equipped with a Micro ToupH electrode (9618S-10D, HORIBA, Ltd.) after a three-point calibration at pH 4.0, 7.0, and 9.0. Following, the concentration of lactate in the homogenates was determined using a multi-assay analyzer (GM7 Micro-Stat; Analox Instruments Ltd., London, UK) after calibration with an 8.0-M standard lactate solution (GMRD-103, Analox Instruments). A 20-µL aliquot of centrifuged supernatant was used for the measurement. In the *in vitro* experiments, lactate concentrations of the cultured cells and culture supernatant were measured using the lactate colorimetric assay kit II (K627-100; BioVision, Milpitas, CA, USA). Light transmittance was measured at a wavelength of 450 nm using a spectrophotometer (DU730; Beckman Coulter, Tokyo, Japan).

### Statistical analysis

The data were analyzed by Student’s t-test, one-way analysis of variance (ANOVA), two-way ANOVA, followed by Tukey’s honestly significant difference post-hoc test, or linear regression analysis using R version 3.5.2 or SAS Studio software version 9.4 (SAS Institute Inc., Cary, NC, USA).

## Data availability

The original blot images are provided in Extended data figures. The full statistical results are provided in Extended data tables.

## Acknowledgements

We thank Giovanni Sala for invaluable comments on statistics. We also thank Tamaki Murakami, Chikako Ozeki, Wakako Hasegawa, Yumiko Mobayashi, Yoko Kagami, Harumi Mitsuya, Yoshihiro Takamiya, Satoko Hattori and Miho Tanaka for their technical assistance. This work was supported by MEXT KAKENHI Grant Number JP16H06462, JP17H06417, and JP17H06413, MEXT Promotion of Distinctive Joint Research Center Program Grant Number JPMXP0618217663, JSPS KAKENHI Grant Number JP20H00522, JP18K07378, JP19K06502, and JP16H06276 (AdAMS), and AMED Strategic Research Program for Brain Sciences Grant Number JP18dm0107101.

## Author information

### Contributions

H.H. and T.M. designed experiments. H.H. wrote the manuscript. K.N., K.K., and T.M. helped draft the manuscript. H.H., H.S., H.O., and A.T. performed the behavioral tests and analyzed the data. K. K. and M. N. performed STED microscopy analysis. T.M. supervised all aspects of the present study. All authors have approved the final manuscript.

### Corresponding authors

Correspondence to Tsuyoshi Miyakawa.

## Ethics declarations

The authors declare no competing interests.

