## Supplemental Figures for "Protein lactylation induced by neural excitation"

**Full title:** Protein lactylation induced by neural excitation

**Short title:** Protein lactylation in the brain

**Extended data figures and tables**

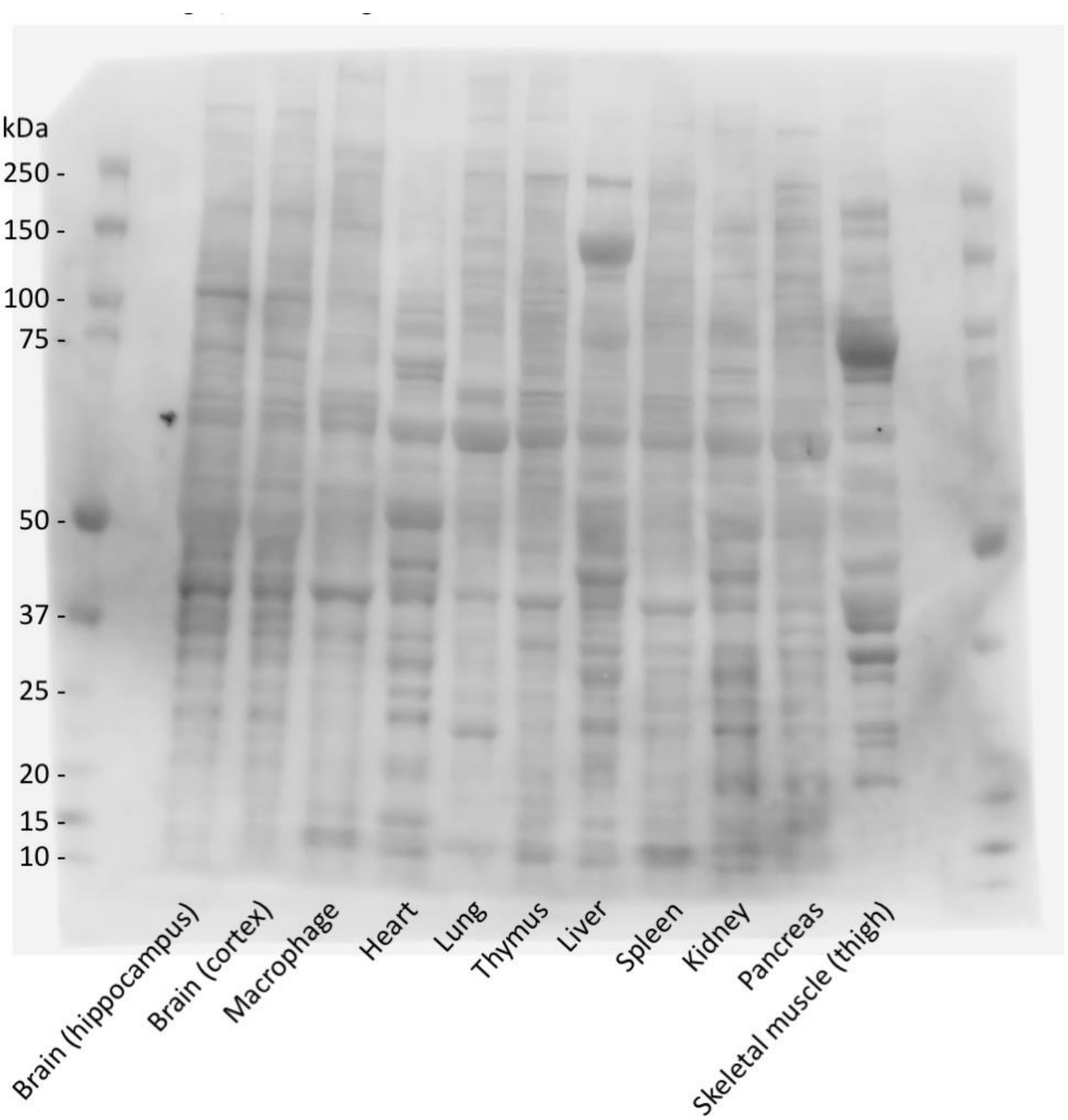


**Extended Data Fig. 1, related to Fig. 1a.** Original blot of Kla in the brain regions and peripheral tissues of mice.

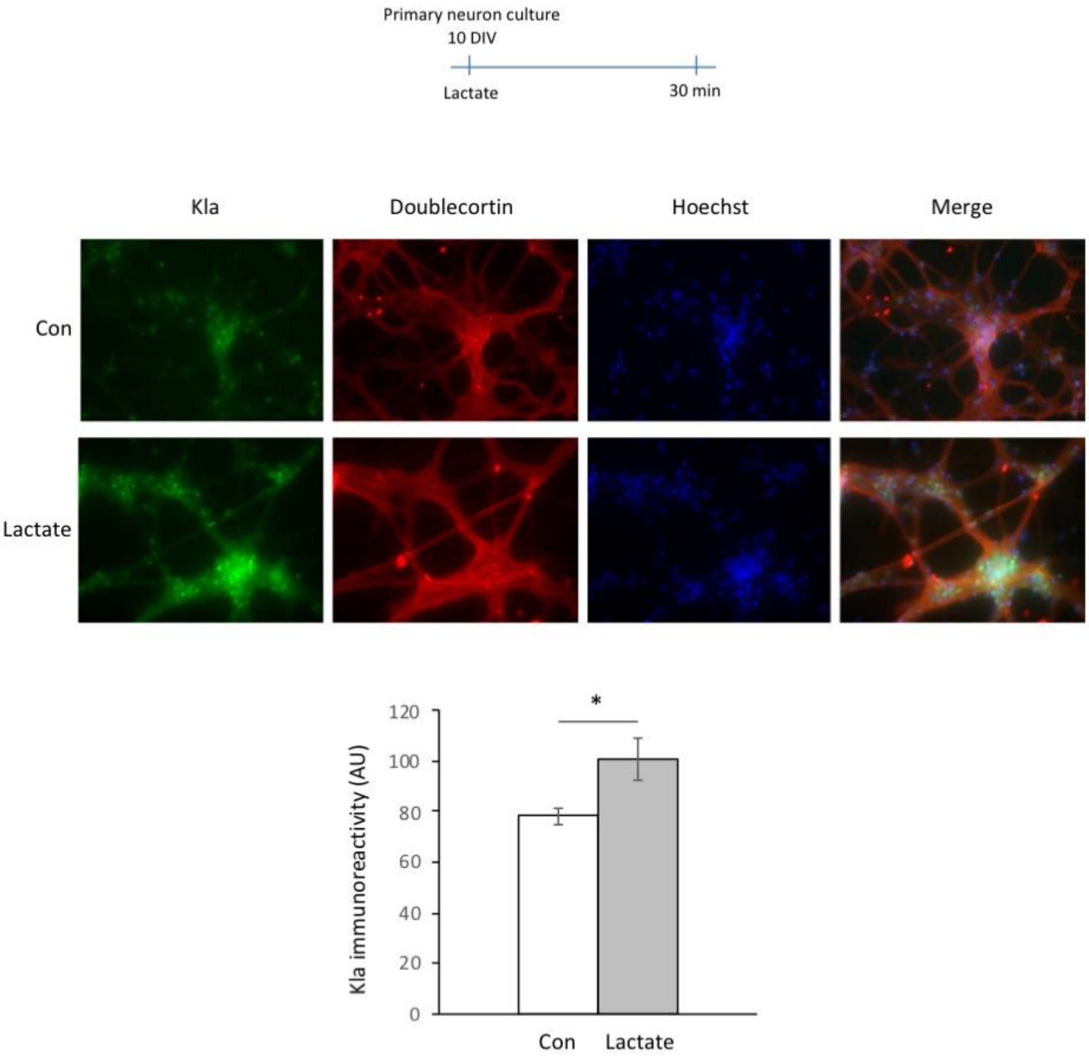


**Extended Data Fig. 2, related to Fig. 1d.** Kla immunostaining of the 10 DIV hippocampal neurons treated with or without 20 mM lactate. Neurons are identified by double staining with doublecortin antibody. N = 3 replicates for each group; *P = 0.010, Students t-test.

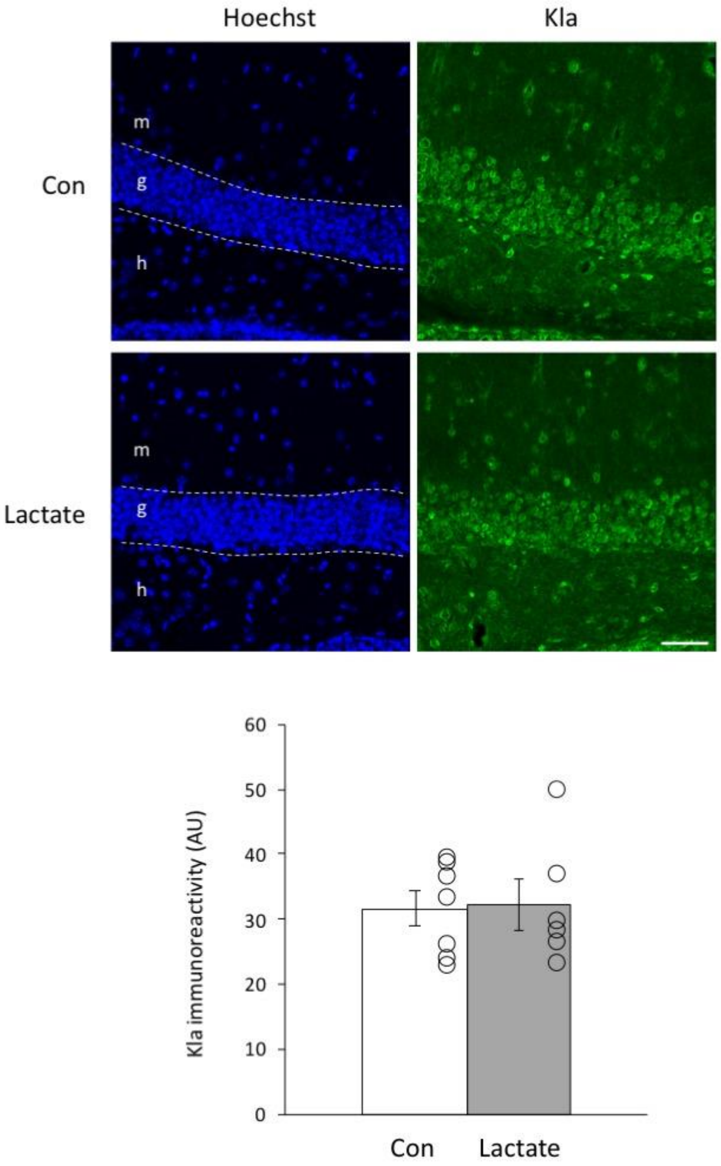


**Extended Data Fig. 3, related to Fig. 1f–h.** Kla immunostaining in the mouse hippocampal dentate gyrus chronically treated with lactate (1 g/kg/day for 21 days). Immunoreactivity in the granule cell layer is quantified. P = 0.86, Student's t-test; N = 6 and 7 mice, respectively; g: granule cell layer; h: hilus; m: molecular layer; Scale bar: 50 µm.

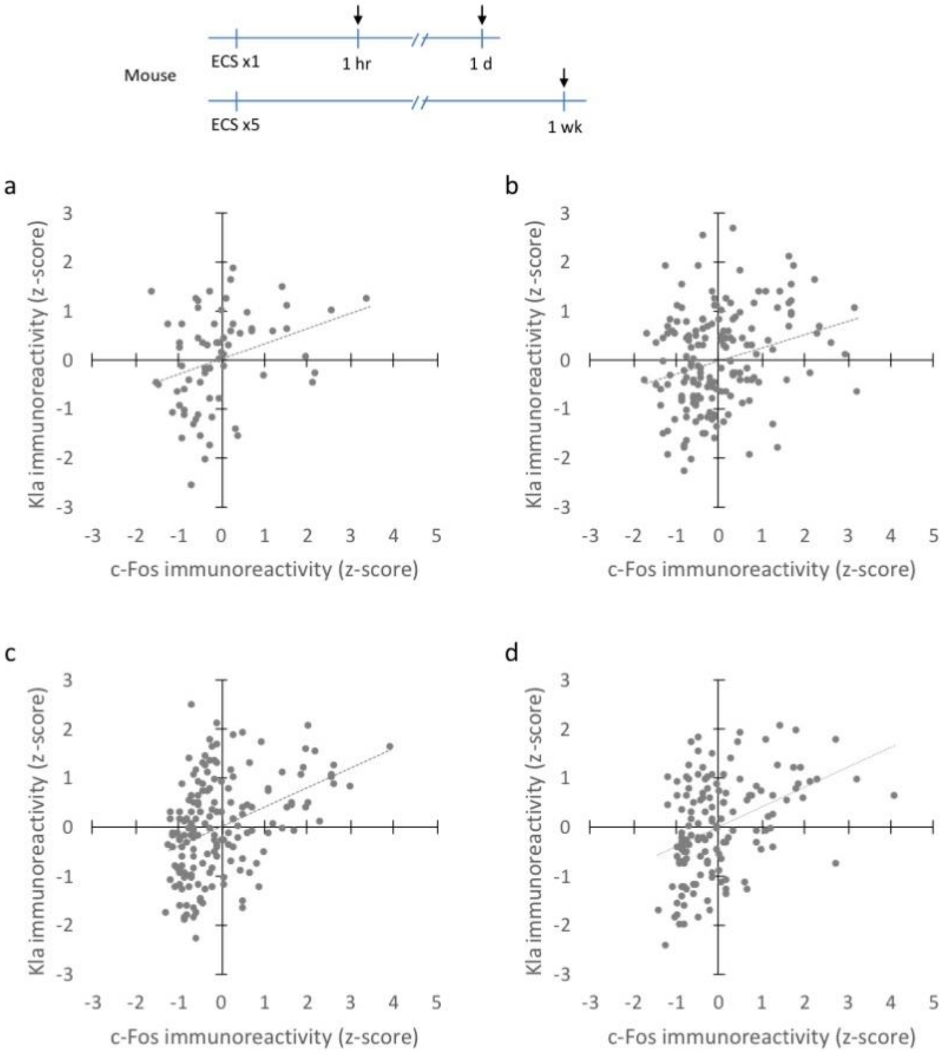


**Extended Data Fig. 4, related to Fig. 2j.** (a–d) Scatterplots showing the correlations between Kla and c-Fos immunoreactivity in individual prefrontal cortex cells in each condition: control (a; r = 0.31, P = 0.0089, N = 69 cells from 3 mice), 1 hour after one session of electroconvulsive stimulation (ECS) (b; r = 0.27, P = 0.00044, N = 164 cells from 4 mice),

1 day after one session of ECS (c; $r = 0.40$, $P = 5.65 \times 10^{-8}$, $N = 169$ cells from 4 mice), and 1 week after five sessions of ECS (once per day for five consecutive days) (d; $r = 0.41$, $P = 5.81 \times 10^{-7}$, $N = 141$ cells from 4 mice). Control mice were handled similarly without ECS treatment and sampled 1 hour after handling. Z-scores are calculated for each variable in each condition.

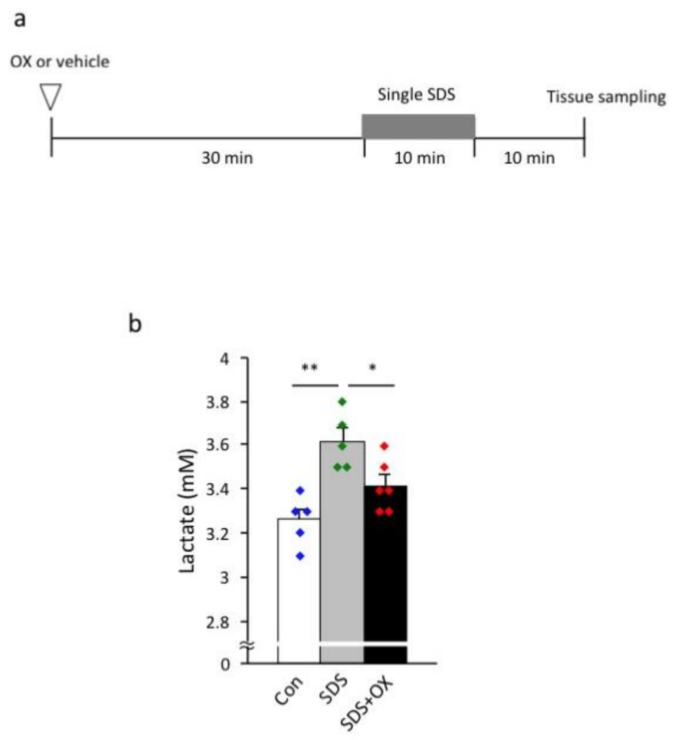


**Extended Data Fig. 5, related to Fig. 3.** (a) Mice were intraperitoneally treated with 1 g/kg sodium oxamate (OX) or vehicle (saline) 30 minutes before being exposed to social defeat stress for 10 minutes. The mice were returned to their home cage and after 10 minutes, brain sampling was performed. (b) Bar graphs showing brain lactate levels in three treatment conditions. *P < 0.05, **P < 0.01, one-way analysis of variance, followed by post-hoc pairwise comparisons. N = 5, 5, and 6 mice, respectively.

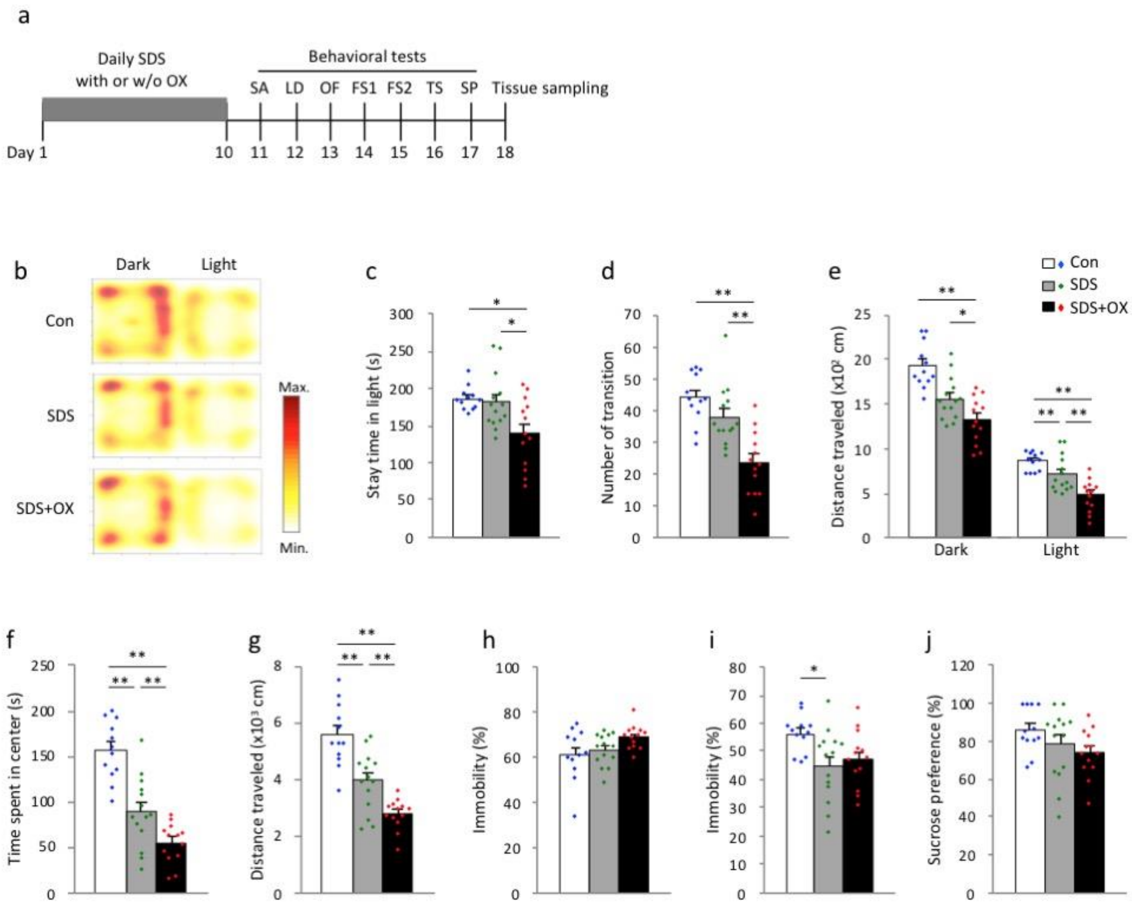


**Extended Data Fig. 6, related to Fig. 3.** Behavioral data of mice exposed to repeated social defeat stress (SDS). (a) Sodium oxamate (OX) or vehicle (saline) was intraperitoneally injected 30 minutes before each SDS session. Control mice were treated with vehicle daily. FS: Porsolt forced swim test; LD: light-dark transition test; OF: open field test; SA: social avoidance test; SP: sucrose preference test; TS: tail suspension test. (b–e) Light/dark transition test. Heat map showing time spent in the dark and light boxes (b), time spent in the light box (c), number of transitions between the dark and light boxes (d), and distance traveled in the dark and light boxes (d). (f, g) Open field test. Time spent in the center area (f) and distance traveled in the field (g). (h) Percentage of Immobility time in the forced swim test. (i) Percentage of immobility time in the tail suspension test. (j) Percentage sucrose preference. Values are means ± SEM. Each dot represents one mouse. *$P < 0.05$, **$P < 0.01$, one-way analysis of variance, followed by Tukey’s post-hoc test; SEM: standard error of the mean.

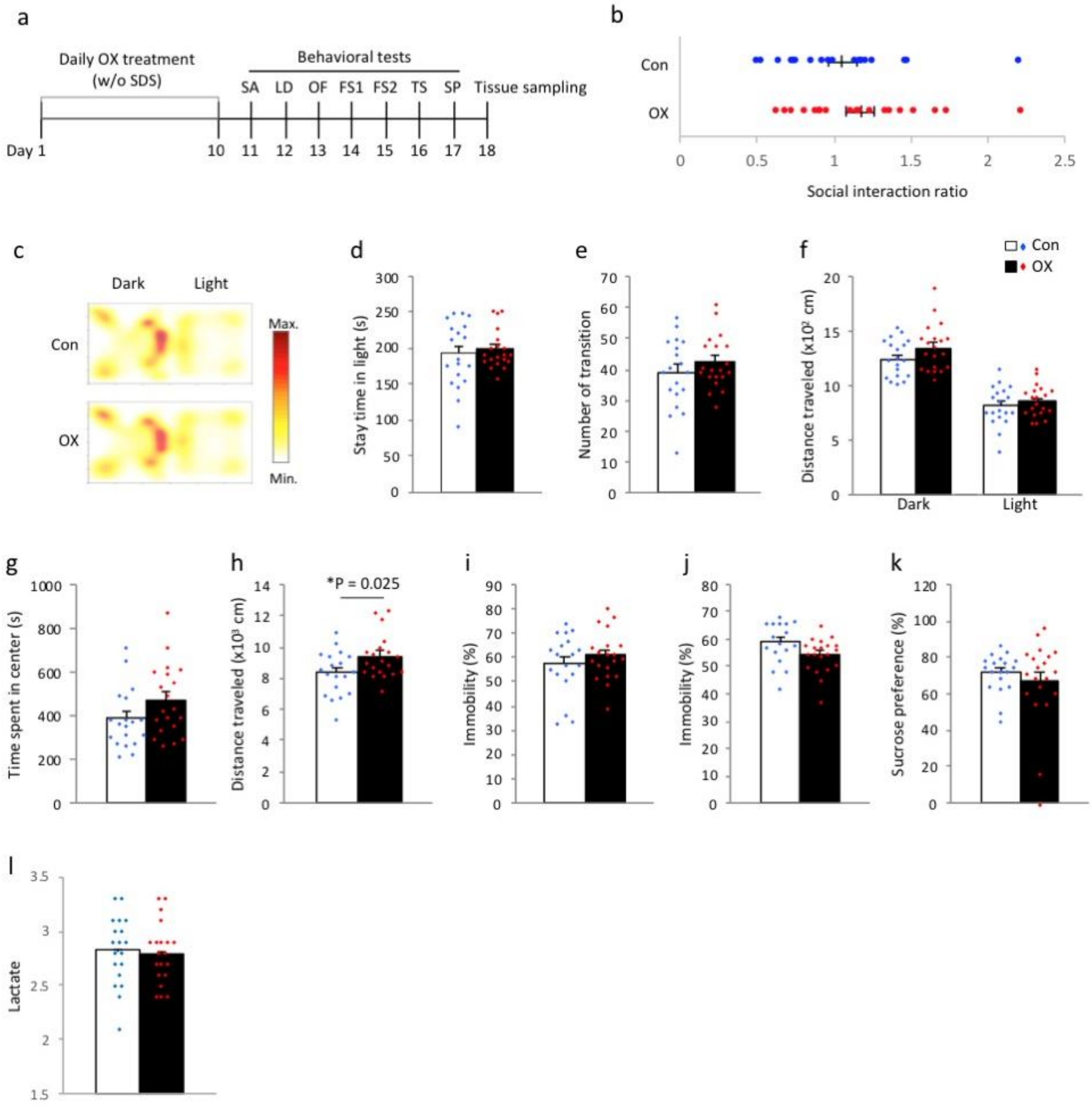


**Extended Data Fig. 7, related to Fig. 3.** Repeated treatment with sodium oxamate does not affect anxiety-like behaviors or brain lactate levels. (a) Timeline of the experiments. SDS: social defeat stress; OX: sodium oxamate; SA: social avoidance test; LD: light/dark transition test; OF: open field test; FS: Porsolt forced swim test; TSP: tail suspension test; SP: sucrose preference test. (b) Social avoidance test. (c–f) Light/dark transition test. Heat map showing (c) the time spent in the dark and light boxes, (d) the time spent in the light box, (e) the number of transitions between the dark and light boxes, and (f) the distance traveled in the dark and light boxes. (g, h) Open field test. (g) Time spent in the center area and (h) distance traveled in the field. (i) Percentage of immobility time in the forced swim test. (j) Percentage of immobility time in the tail suspension test. (k) Percentage sucrose preference. (l) Brain lactate levels. Values are means ± SEM. Each dot represents one mouse.

*P < 0.05, Student’s t-test; SEM: standard error of the mean.

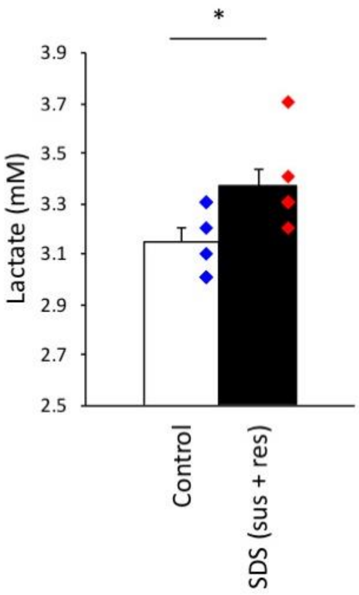


**Extended Data Fig. 8, related to Fig. 3.** The presence of increased lactate levels in the brains of mice exposed to social defeat stress is confirmed by an independent experiment performed at a different institute.

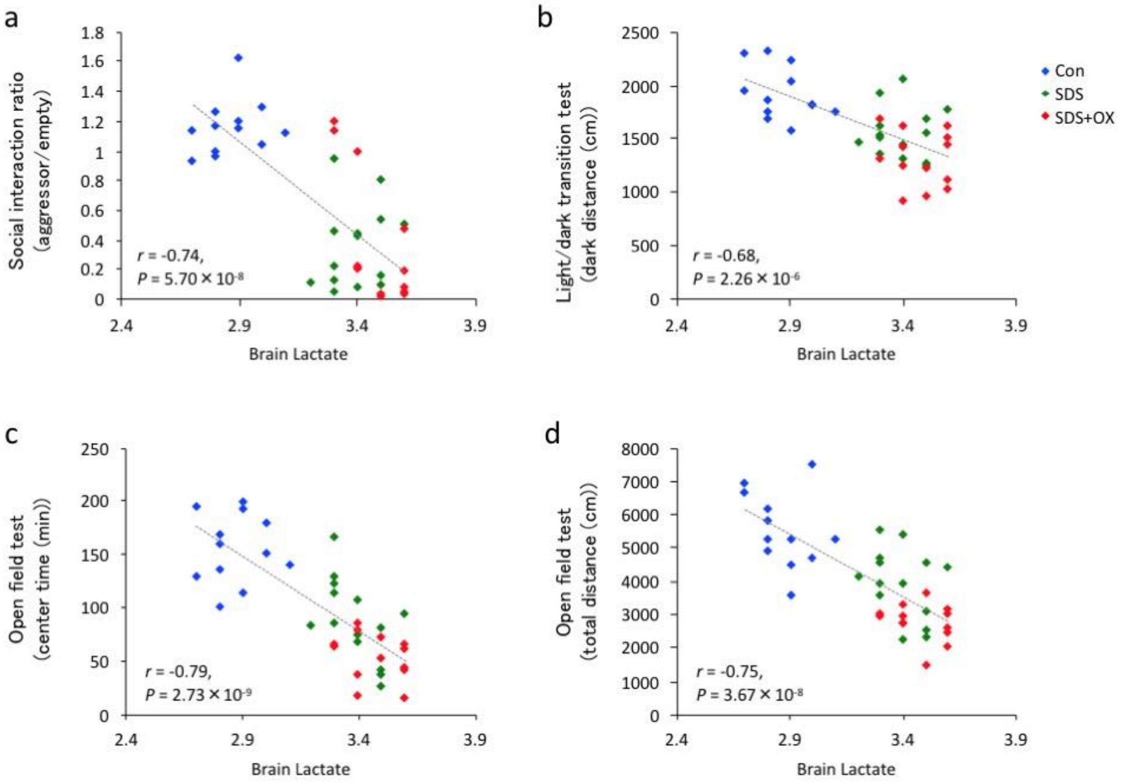


**Extended Data Fig. 9, related to Fig. 3.** Scatterplots showing significant correlations between brain lactate levels and (a) the social interaction ratio measured by the social avoidance test, (b) the distance traveled in the dark box in the light/dark transition test, (c) the time spent in center area in the open field test and (d) the total distance traveled in the open field test.

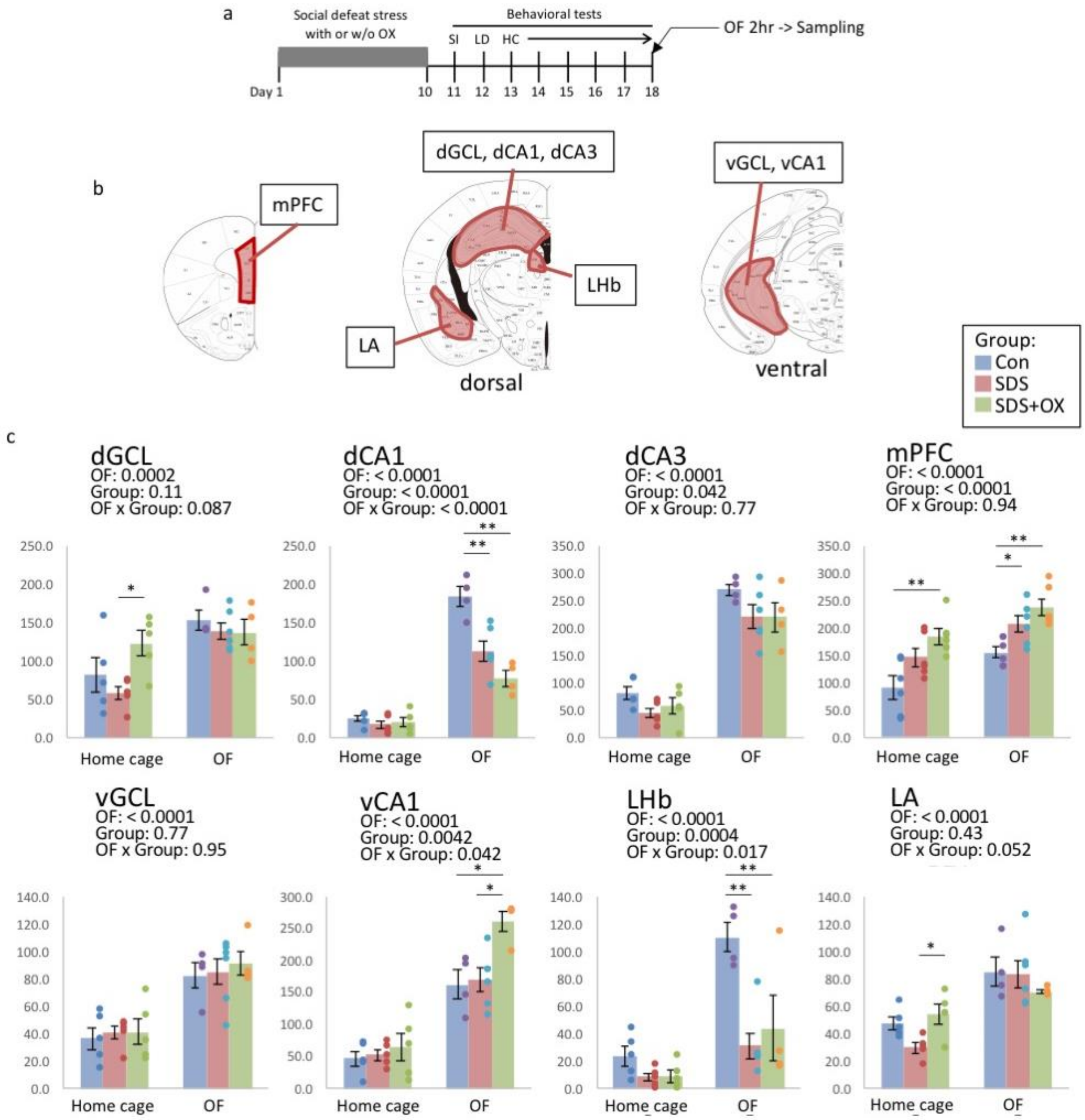


**Extended Data Fig. 10, related to Fig. 3.** (a) Timeline of the experiments. SDS: social defeat stress; OX: sodium oxamate; SA: social avoidance test; LD: light/dark transition test; HC: home cage locomotor activity monitoring test; OF: open field test. (b) Brain areas examined for numbers of c-Fos-positive cells. Adapted from [32]. (c) Bar graphs showing numbers of c-Fos-positive cells in each brain region. *P < 0.05, **P < 0.01, two-way analysis of variance, followed by Tukey's honestly significant difference post-hoc test. dCA1, dorsal CA1; dCA3, dorsal CA3; dGCL, dorsal granule cell layer; LA, lateral amygdala; LHb, lateral habenula; mPFC, medial prefrontal cortex; vGCL, ventral granule cell layer; vCA1, ventral CA1.

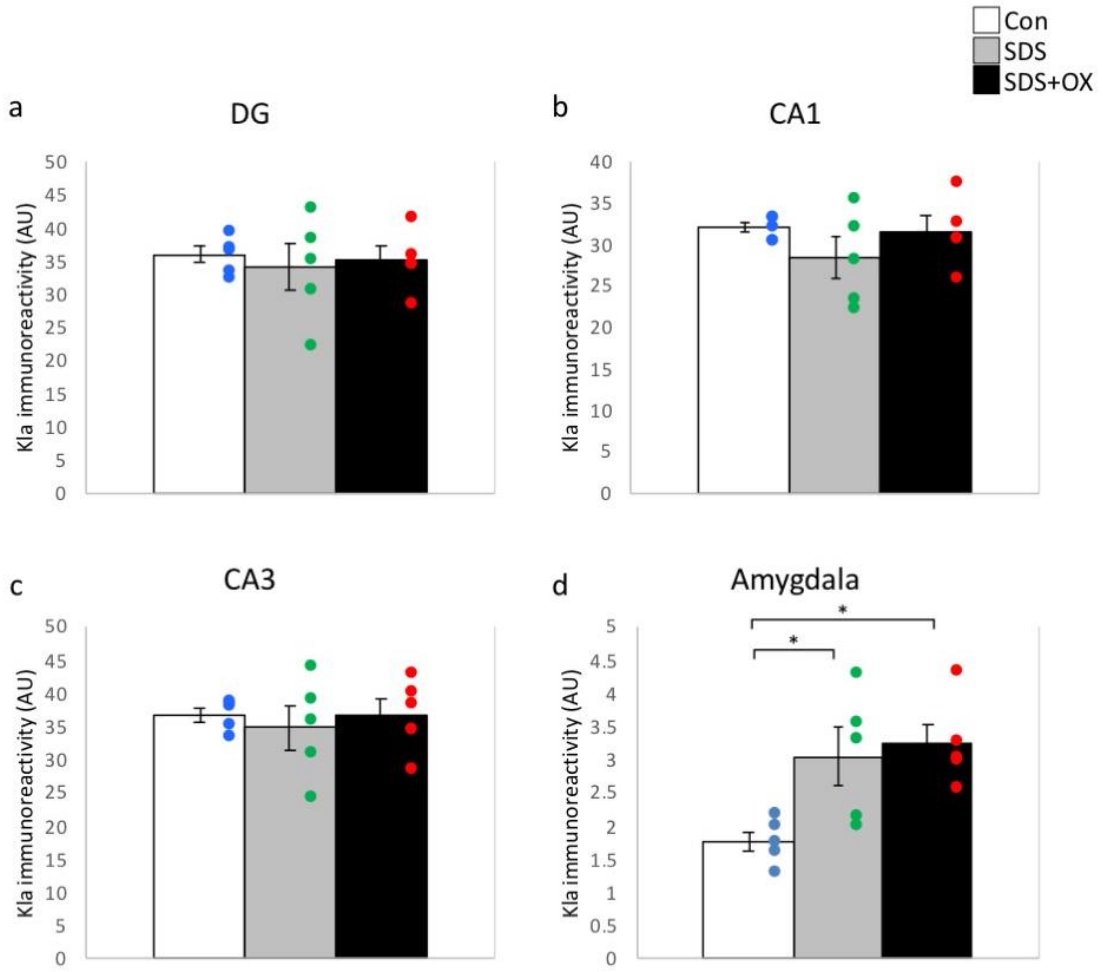


**Extended Data Fig. 11, related to Fig. 3.** Kla immunoreactivity in the dorsal hippocampus (i.e., the granule cell layer of the dentate gyrus (a) and the pyramidal cell layer of the CA1 (b) and CA3 (c) regions) and the basolateral amygdala (d). *$P < 0.05$, **$P < 0.01$, one-way analysis of variance, followed by Tukey's honestly significant difference post-hoc test.

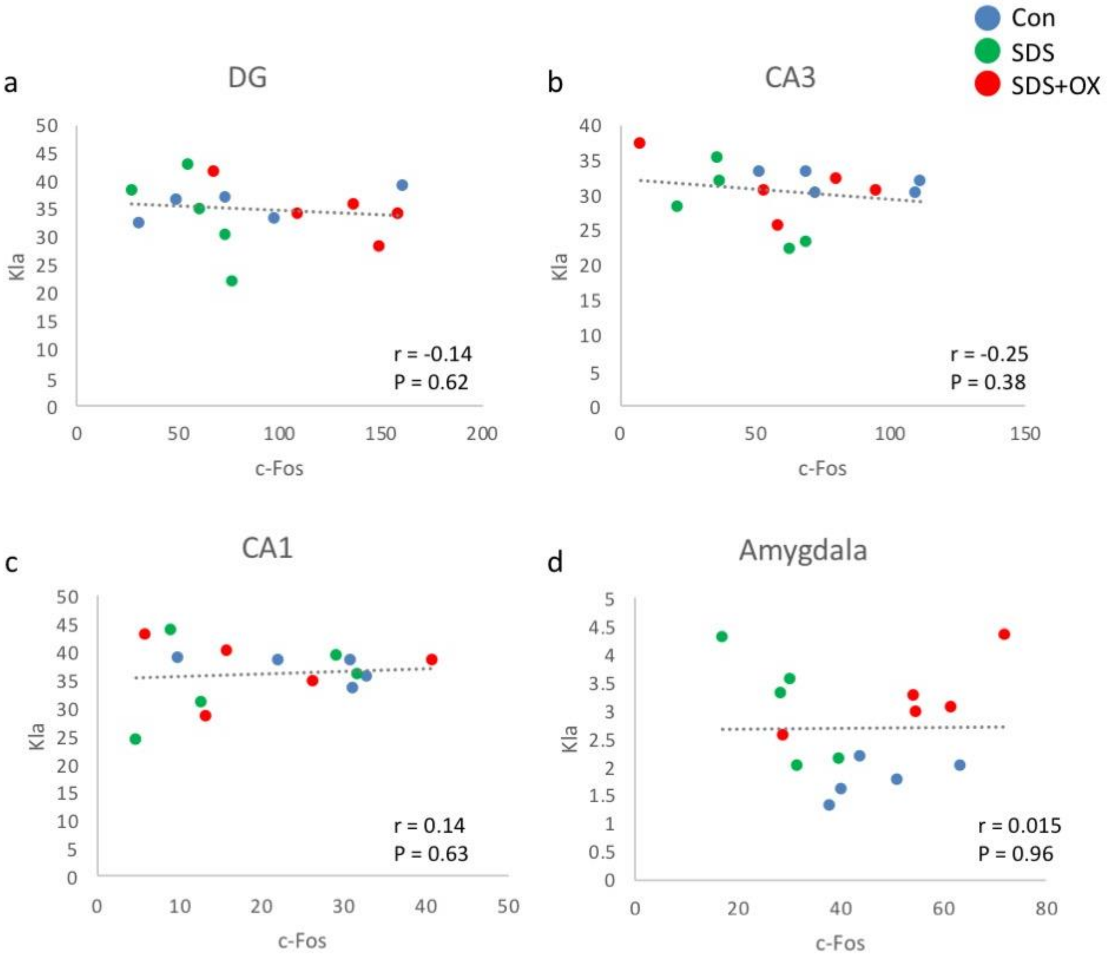


**Extended Data Fig. 12, related to Fig. 3.** Scatterplots showing the correlation between Kla immunoreactivity and numbers of c-Fos-positive cells in the hippocampus (i.e., the granule cell layer of the dentate gyrus (a) and the pyramidal-cell layer of the CA1 (b) and CA3 (c) regions) and the basolateral amygdala (d).

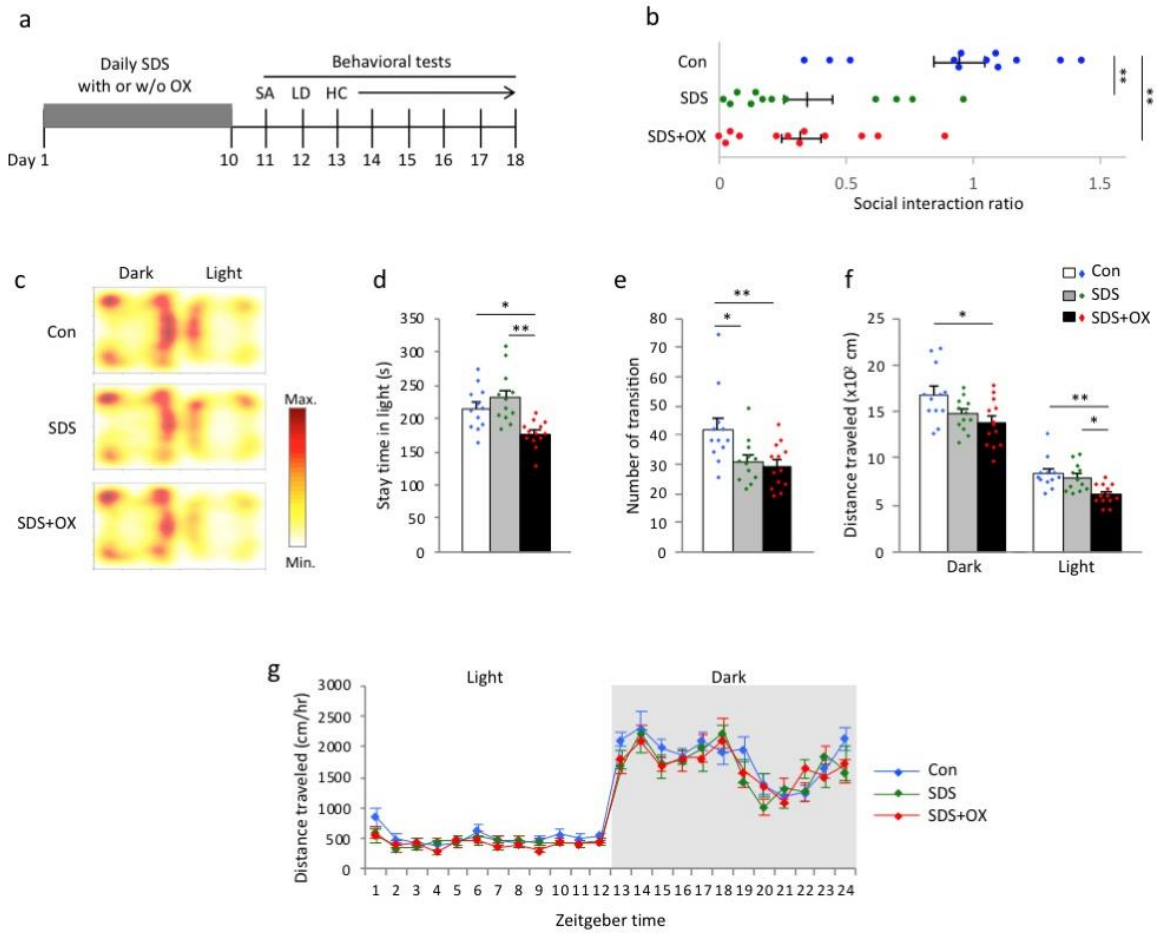


**Extended Data Fig. 13, related to Fig. 3.** Home cage locomotor activity monitoring of mice exposed to repeated social defeat stress (SDS). (a) Sodium oxamate or vehicle (saline) was intraperitoneally injected 30 minutes before each SDS session. Control mice were treated with vehicle daily. HC: home cage locomotor activity monitoring test; LD: light-dark transition test; SA: social avoidance test. (b) Social avoidance test. (c–f) Light/dark transition test. Heat map showing (c) the time spent in the dark and light boxes, (d) the time spent in the light box, (e) the number of transitions between the dark and light boxes, and (f) the distance traveled in the dark and light boxes. (g) Results of monitoring home-cage locomotor activity. $P = 0.17$, repeated measures analysis of variance.

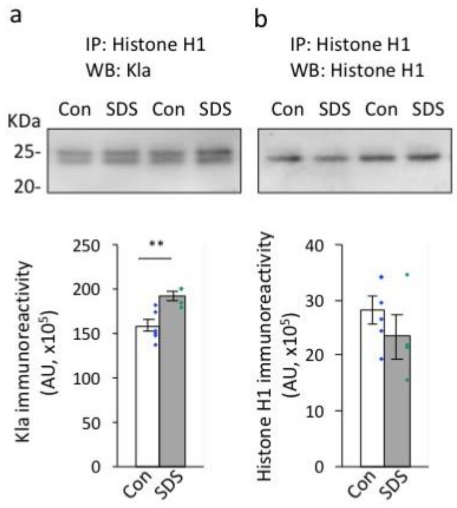


**Extended Data Fig. 14, related to Fig. 4.** (a–b) Immunoprecipitation of mouse prefrontal cortex (PFC) with histone H1 antibody and subsequent western blot analysis with Kla antibody (a) and histone H1 antibody (b). Bar graphs show immunoreactivity for Kla (a) and histone H1 (b) at approximately 25 kDa.

**Extended Data Table 1.** Multiple regression analysis

**Extended Data Table 2.** Candidate lysine-lactylated proteins in the mouse prefrontal cortex

**Extended Data Table 3.** GO enrichment analysis

**Extended Data Table 4–12.** Full statistical results
